# Music-selective neural populations arise without musical training

**DOI:** 10.1101/2020.01.10.902189

**Authors:** Dana Boebinger, Sam Norman-Haignere, Josh McDermott, Nancy Kanwisher

## Abstract

Recent work has shown that human auditory cortex contains neural populations anterior and posterior to primary auditory cortex that respond selectively to music. However, it is unknown how this selectivity for music arises. To test whether musical training is necessary, we measured fMRI responses to 192 natural sounds in 10 people with almost no musical training. When voxel responses were decomposed into underlying components, this group exhibited a music-selective component that was very similar in response profile and anatomical distribution to that previously seen in individuals with moderate musical training. We also found that musical genres that were less familiar to our participants (e.g., Balinese *gamelan*) produced strong responses within the music component, as did drum clips with rhythm but little melody, suggesting that these neural populations are broadly responsive to music as a whole. Our findings demonstrate that the signature properties of neural music selectivity do not require musical training to develop, showing that the music-selective neural populations are a fundamental and widespread property of the human brain.

**NEW & NOTEWORTHY:** We show that music-selective neural populations are clearly present in people without musical training, demonstrating that they are a fundamental and widespread property of the human brain. Additionally, we show music-selective neural populations respond strongly to music from unfamiliar genres as well as music with rhythm but little pitch information, suggesting that they are broadly responsive to music as a whole.

## INTRODUCTION

Music is uniquely and universally human (Mehr et al., 2019) and musical abilities arise early in development (Trehub, 2003). Recent evidence has revealed neural populations in bilateral non-primary auditory cortex that respond selectively to music and thus seem likely to figure importantly in musical perception and behavior (Leaver and Rauschecker, 2010; Rogalsky et al., 2011; Fedorenko et al., 2012; Tierney et al., 2013; LaCroix et al., 2015; Norman-Haignere et al., 2015, 2019). How does such selectivity arise? Most members of Western societies have received at least some explicit musical training in the form of lessons or classes, and it is possible that this training leads to the emergence of music-selective neural populations. However, most Western individuals, including non-musicians, also implicitly acquire knowledge of musical structure from a lifetime of exposure to music (Bigand, 1983; Bigand and Pineau, 1997; Koelsch et al., 2000; Tillmann et al., 2000; Tillmann, 2005; Bigand and Poulin-Charronnat, 2006). Thus, another possibility is that this type of passive experience with music is sufficient for the development of cortical music selectivity. The roles of these two forms of musical experience in the neural representation of music are not understood. Here, we directly test whether explicit musical training is necessary for the development of music-selective neural responses, by testing whether music-selective responses are robustly present – with similar response characteristics and anatomical distribution – in individuals with little or no explicit training.

Why might explicit musical training be necessary for neural tuning to music? The closest analogy in the visual domain is learning to read, where several studies have shown that selectivity to visual orthography (Baker et al., 2007) arises in high-level visual cortex only after children are taught to read (Dehaene et al., 2010; Dehaene-Lambertz et al., 2018). In audition, exposure to specific sounds can elicit long-term changes in auditory cortex, such as sharper tuning of individual neurons (Recanzone et al., 1993; Fritz et al., 2003; Lee and Middlebrooks, 2011) and expansion of cortical maps (Recanzone et al., 1993; Polley et al., 2006; Bieszczad and Weinberger, 2010). These changes occur primarily for behaviorally relevant stimulus features (Ahissar et al., 1992, 1998; Fritz et al., 2005; Ohl and Scheich, 2005; Polley et al., 2006) related to the intrinsic reward value of the stimulus (Bakin and Weinberger, 1996; Fritz et al., 2005; David et al., 2012), and thus are closely linked to the neuromodulatory system (Bao et al., 2001; Kilgard et al., 2001; Blake et al., 2006). Additionally, the extent of cortical map expansion is correlated with the animal’s subsequent improvement in behavioral performance (Recanzone et al., 1993; Rutkowski and Weinberger, 2005; Polley et al., 2006; Bieszczad and Weinberger, 2010, 2012; Reed et al., 2011). Most of this prior work on experience-driven plasticity in auditory cortex has been done in animals undergoing extensive training on simple sensory sound dimensions, and it has remained unclear how the results from this work might generalize to humans in more natural settings with higher-level perceptual features. Musical training in humans provides a unique way to investigate this question, as it meets virtually all of these criteria for eliciting functional plasticity: playing music requires focused attention, fine-grained sensory-motor coordination, it is known to engage the neuromodulatory system (Blood and Zatorre, 2001; Salimpoor et al., 2011, 2013), and expert musicians often begin training at a young age and hone their skills over many years.

Although many prior studies have measured neural changes as a result of auditory experience (Golestani et al., 2011; Teki et al. 2012), including comparing responses in musicians and non-musicians (Ohnishi et al., 2001; Pantev et al., 2001; Shahin et al., 2003; Fujioka et al., 2004, 2005; Besson et al., 2007; Wong et al., 2007; Dick et al., 2011; Lee and Noppeney, 2011; Ellis et al., 2012, 2013; Angulo-Perkins et al., 2014; Doelling and Poeppel, 2015; Lappe et al., 2016), it remains unclear whether any tuning properties of auditory cortex depend on musical training. Previous studies have found that fMRI responses to music are larger in musicians compared to non-musicians in posterior superior temporal gyrus (Ohnishi et al., 2001; Dick et al., 2011; Angulo-Perkins et al., 2014). However, these responses were not shown to be selective for music, leaving the relationship between musical training and cortical music selectivity unclear.

Music selectivity is weak when measured in raw voxel responses using standard voxel-wise fMRI analyses, due to spatial overlap between music-selective neural populations and neural populations with other selectivities (e.g. pitch). To overcome these challenges, Norman-Haignere et al. (2015) used voxel decomposition to infer a small number of component response profiles that collectively explained voxel responses throughout auditory cortex to large set of natural sounds. This approach makes it possible to disentangle the responses of neural populations that overlap within voxels, and has previously revealed a neural population with clear selectivity for music compared to both other real-world sounds (Norman-Haignere et al., 2015) and synthetic control stimuli matched to music in many acoustic properties (Norman-Haignere and McDermott, 2018). These results have recently been confirmed by intracranial recordings, which show individual electrodes with clear selectivity for music (Norman-Haignere et al., 2019). Although Norman-Haignere et al. (2015) did not include actively practicing musicians, many of the participants had substantial musical training earlier in their lives.

Here, we test whether music selectivity arises only after explicit musical training. To this end, we probed for music selectivity in people with almost no musical training. On the one hand, if explicit musical training is necessary for the existence of music-selective neural populations, music selectivity should be weak or absent in these non-musicians. If, however, music selectivity does not require explicit training but rather is either innate or arises as a consequence of passive exposure to music, then we would expect to see robust music selectivity even in the non-musicians. A group of highly trained musicians was also included for comparison. Using these same methods, we were also able to test whether the inferred music-selective neural population responds strongly to less familiar musical genres (e.g. Balinese *gamelan*), and to drum clips with rich rhythm but little melody.

Note that this is not a traditional group comparison study contrasting musicians and non-musicians in an attempt to ascertain whether musical training has any detectable effect on music selective neural responses, as it would be unrealistic to collect the amount of data that would be necessary for a direct comparison between groups (see “Direct group comparisons of music selectivity” in the **Supplemental Information**). Rather, our goal was to ask whether the key properties of music selectivity described in our earlier study are present in each group when analyzed separately, thus determining whether or not explicit training is necessary for the emergence of music selective responses in the human brain.

## MATERIALS & METHODS

### Participants

Twenty young adults (14 female, mean = 24.7 years, SD = 3.8 years) participated in the experiment: 10 non-musicians (6 female, mean = 25.8 years, SD = 4.1 years) and 10 musicians (8 female, mean = 23.5 years, SD = 3.3 years). This number of participants was chosen because our previous study (Norman-Haignere et al., 2015) was able to infer a music-selective component from an analysis of ten participants. Although these participants were described as “non-musicians” (defined as no formal training in the five years preceding the study) many of the participants had substantial musical training earlier in life. We therefore used stricter inclusion criteria to recruit 10 musicians and 10 non-musicians for the current study, in order to have statistical power within each group comparable to that of our previous study.

To be classified as a non-musician, participants were required to have less than two years of total music training, which could not have occurred either before the age of seven or within the last five years. Out of the ten non-musicians in our sample, eight had zero years of musical training, one had a single year of musical training (at the age of 20), and one had two years of training (starting at age 10). These training measures do not include any informal “music classes” included in participants’ required elementary school curriculum, because (at least in the United States) these classes are typically compulsory, are only for a small amount of time per week (e.g. 1 hour), and primarily consist of simple sing-a-longs. Inclusion criteria for musicians were beginning formal training before the age of seven (Penhune, 2011), and continuing training until the current day. Our sample of ten musicians had an average of 16.30 years of training (ranging from 11-23 years, SD = 2.52). Details of participants’ musical experience can be found in **Table 1**.

**Table 1.** Details of participants’ musical backgrounds and training, as measured by a self-report questionnaire.

|  | Subject # | Instrument | Age of Onset (Years) | Years of Lessons | Years of Regular Practice | Years of Training | Hours of Weekly Practice | Hours of Daily Music Listening |
| --- | --- | --- | --- | --- | --- | --- | --- | --- |
| NON-MUSICIANS | 1 |  |  | 0 | 0 | 0 | 0 | 0 |
|  | 2 |  | 20 | 1 | 0 | 1 | 0 | 2 |
|  | 3 |  |  | 0 | 0 | 0 | 0 | 0.25 |
|  | 4 |  |  | 0 | 0 | 0 | 0 | 5 |
|  | 5 |  |  | 0 | 0 | 0 | 0 | 1 |
|  | 6 |  |  | 0 | 0 | 0 | 0 | 1.5 |
|  | 7 |  | 10 | 1.5 | 0 | 1.5 | 0 | 1 |
|  | 8 |  |  | 0 | 0 | 0 | 0 | 0.1 |
|  | 9 |  |  | 0 | 0 | 0 | 0 | 0.5 |
|  | 10 |  |  | 0 | 0 | 0 | 0 | 5 |
|  | mean |  | 24.4 | 0.3 | 0 | 0.3 | 0 | 1.64 |
| MUSICIANS | 11 | violin | 7 | 11 | 11 | 11 | 2 | 2 |
|  | 12 | piano, French horn | 5 | 7 | 18 | 12 | 6 | 4 |
|  | 13 | piano, bass | 6 | 12 | 18 | 18 | 2 | 2 |
|  | 14 | piano | 3 | 15 | 15 | 15 | 2 | 2.5 |
|  | 15 | piano, violin, viola | 5 | 17 | 19 | 20 | 3 | 4 |
|  | 16 | piano, flute | 5 | 13 | 13 | 13 | 12 | 3 |
|  | 17 | flute | 7 | 12 | 13 | 15 | 15 | 3 |
|  | 18 | piano, cello, flute | 4 | 19 | 15 | 18 | 25 | 1 |
|  | 19 | violin | 3 | 18 | 18 | 18 | 4 | 10 |
|  | 20 | violin, clarinet | 4 | 13 | 8 | 23 | 4 | 5 |
|  | mean |  | 5.2 | 13.7 | 14.8 | 16.3 | 7.1 | 3.65 |
|  | grand mean |  | 14.8 | 7 | 7.4 | 8.3 | 3.55 | 2.64 |

Non-parametric Wilcoxon rank sum tests indicated that there were no significant group differences in median age (musician median = 24.0 years, SD = 3.3, non-musician median = 25.0 years, SD = 4.1, z = −1.03, p = 0.30, effect size r = −0.23), post-secondary education (i.e. formal education after high school; musician median = 6.0 years, SD = 8.2 years, non-musician median = 6.5 years, SD = 7.5 years, z = −0.08, p = 0.94, effect size r = −0.02), or socioeconomic status as measured by the Barrett Simplified Measure of Social Status questionnaire (BSMSS; (Barratt, 2006) (musician median = 54.8, SD = 7.1, non-musician median = 53.6, SD = 15.4, z = 0.30, p = 0.76, effect size r = 0.07). Note that we report the group standard deviations because this measure is more robust than the interquartile range with our modest sample size of 10 participants per group. All participants were native English speakers and had normal hearing (audiometric thresholds <25 dB HL for octave frequencies 250Hz to 8kHz). The study was approved by MIT’s human participants review committee (COUHES), and written informed consent was obtained from all participants.

To validate participants’ self-reported musicianship, we measured participants’ abilities on a variety of psychoacoustical tasks for which prior evidence suggested that musicians would outperform non-musicians, including frequency discrimination, sensorimotor synchronization, melody discrimination, and “sour note” detection. As predicted, musician participants outperformed non-musician participants on all behavioral psychoacoustic tasks. See **Supplemental Information** for more details.

### Study design

Each participant underwent a 2-hour behavioral testing session as well as three 2-hour fMRI scanning sessions. During the behavioral session, participants completed an audiogram to rule out the possibility of hearing loss, filled out questionnaires about their musical experience, and completed a series of basic psychoacoustic tasks described in **Supplemental Information**.

### Natural sound stimuli for fMRI experiment

Stimuli consisted of 2-second clips of 192 natural sounds. These sounds included the 165-sound stimulus set used in Norman-Haignere et al. (2015), which broadly sample frequently heard and recognizable sounds from everyday life. Examples can be seen in **Figure 1A**. This set of 165 sounds was supplemented with 27 additional music and drumming clips from a variety of musical cultures, so that we could examine responses to rhythmic features of music, as well as comparing responses to more versus less familiar musical genres. Stimuli were ramped on and off with a 25ms linear ramp. During scanning, auditory stimuli were presented over MR-compatible earphones (Sensimetrics S14) at 75 dB SPL.

**Figure 1.**
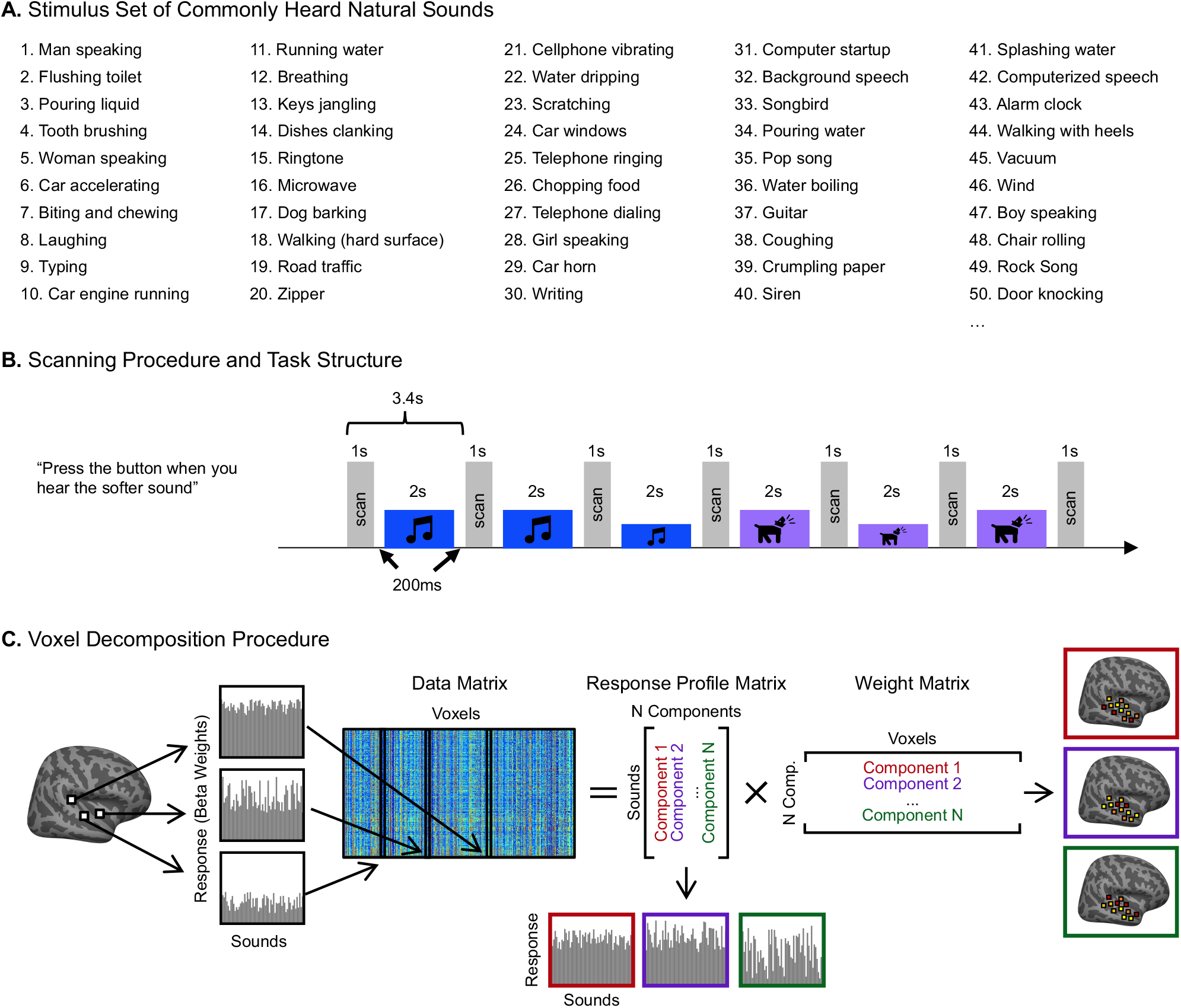
Experimental design and voxel decomposition method. (**A**) Fifty examples from the original set of 165 natural sounds used in Norman-Haignere et al. (2015) and in the current study, ordered by how often participants’ reported hearing them in daily life. An additional 27 music stimuli were added to this set of 165 for the current experiment. (**B**) Scanning paradigm and task structure. Each 2-second sound stimulus was repeated three times consecutively, with one repetition (the second or third) being 12 dB quieter. Subjects were instructed to press a button when they detected this quieter sound. A sparse scanning sequence was used, in which one fMRI volume was acquired in the silent period between stimuli. (**C**) Diagram depicting the voxel decomposition method, reproduced from Norman-Haignere et al. (2015). The average response of each voxel to the 192 sounds is represented as a vector, and the response vector for every voxel from all 20 subjects is concatenated into a matrix (192 sounds x 26,792 voxels). This matrix is then factorized into a response profile matrix (192 sounds x N components) and a voxel weight matrix (N components x 26,792 voxels).

An online experiment (via Amazon’s Mechanical Turk) was used to assign a semantic category to each stimulus, in which 180 participants (95 females; mean age = 38.8 years, SD = 11.9 years) categorized each stimulus into one of fourteen different categories. The categories were taken from Norman-Haignere et al. (2015), with three additional categories (“non-Western instrumental music,” “non-Western vocal music,” “drums”) added to accommodate the additional music stimuli used in this experiment.

A second Amazon Mechanical Turk experiment was run to ensure that American listeners were indeed less familiar with the non-Western music stimuli chosen for this experiment, but that they still perceived the stimuli as “music.” In this experiment, 188 participants (75 females; mean age = 36.6 years, SD = 10.5 years) listened to each of the 62 music stimuli and rated them based on (1) how “musical” they sounded, (2) how “familiar” they sounded, (3) how much they “liked” the stimulus, and (4) how “foreign” they sounded.

### fMRI data acquisition and preprocessing

Similar to the design of Norman-Haignere et al (2015), sounds were presented during scanning in a “mini-block design,” in which each 2-second natural sound was repeated three times in a row. Sounds were repeated because we have found this makes it easier to detect reliable hemodynamic signals. We used fewer repetitions than in our prior study (3 vs. 5), because we wanted to test a larger number of sounds and because we observed similarly reliable responses using fewer repetitions in pilot experiments. Each stimulus was presented in silence, with a single fMRI volume collected between each repetition (i.e. “sparse scanning,” (Hall et al., 1999)). To encourage participants to pay attention to the sounds, either the second or third repetition in each “mini-block” was 12dB quieter (presented at 67 dB SPL), and participants were instructed to press a button when they heard this quieter sound (**Figure 1B**). Overall, participants performed well on this task (musicians: mean = 92.06%, SD = 5.47%; non-musicians: mean = 91.47%, SD = 5.83%; no participant’s average performance across runs fell below 80%). Each of the three scanning sessions consisted of sixteen 5.5-minute runs, for a total of 48 functional runs per participant. Each run consisted of 24 stimulus mini-blocks and five silent blocks during which no sounds were presented. These silent blocks were the same duration as the stimulus mini-blocks, and were distributed evenly throughout each run, providing a baseline. Each specific stimulus was presented in two mini-blocks per scanning session, for a total of six mini-block repetitions per stimulus over the three scanning sessions. Stimulus order was randomly permuted across runs and across participants.

MRI data were collected at the Athinoula A. Martinos Imaging Center of the McGovern Institute for Brain Research at MIT, on a 3T Siemens Prisma with a 32-channel head coil. Each volume acquisition lasted 1 second, and the 2-second stimuli were presented during periods of silence between each acquisition, with a 200ms buffer of silence before and after stimulus presentation. As a consequence, one brain volume was collected every 3.4 seconds (1 second + 2 seconds + 0.2*2 seconds) (TR = 3.4s, TA = 1.02s, TE = 33ms, 90 degree flip angle, 4 discarded initial acquisitions). Each functional acquisition consisted of 48 roughly axial slices (oriented parallel to the anterior-posterior commissure line) covering the whole brain, each slice being 3mm thick and having an in-plane resolution of 2.1 × 2.1mm (96 × 96 matrix, 0.3mm slice gap). An SMS acceleration factor of 4 was used in order to minimize acquisition time (TA = 1.02s). To localize functional activity, a high-resolution anatomical T1-weighted image was obtained for every participant (TR = 2.53 seconds, voxel size: 1mm^3^, 176 slices, 256 × 256 matrix).

Preprocessing and data analysis were performed using FSL software and custom Matlab scripts. Functional volumes were motion-corrected, slice-time-corrected, skull-stripped, linearly detrended, and aligned to each participant’s anatomical image (using FLIRT and BBRegister; (Jenkinson and Smith, 2001; Greve and Fischl, 2009). Motion correction and function-to-anatomical registration was done separately for each run. Preprocessed data were then resampled to the cortical surface reconstruction computed by FreeSurfer (Dale et al., 1999), and smoothed on the surface using a 3mm FWHM kernel to improve SNR. The data were then downsampled to a 2mm isotropic grid on the FreeSurfer-flattened cortical surface.

Next, we estimated the response to each of the 192 stimuli using a general linear model (GLM). Each stimulus mini-block was modeled as a boxcar function convolved with a canonical hemodynamic response function (HRF). The model also included six motion regressors and a first-order polynomial noise regressor to account for linear drift in the baseline signal. Note that this analysis differs from our prior paper (Norman-Haignere et al., 2015), in which signal averaging was used in place of a GLM. We made this change because BOLD responses were estimated more reliably using an HRF-based GLM, potentially due to the use of shorter stimulus blocks causing more overlap between BOLD responses to different stimuli.

### Voxel selection

The first step of this analysis method is to determine which voxels serve as input to the decomposition algorithm. All analyses were carried out on voxels within a large anatomical constraint region encompassing bilateral superior temporal and posterior parietal cortex (**Supplemental Figure S1**), as in Norman-Haignere et al. (2015). In practice, the vast majority of voxels with a robust and reliable response to sound fell within this region (**Supplemental Figure S2**), which explains why our results were very similar with and without this anatomical constraint (**Supplemental Figure S3**). Within this large anatomical region, we selected voxels that displayed a significant (p < 0.001, uncorrected) response to sound (pooling over all sounds compared to silence). This consisted of 51.45% of the total number of voxels within our large constraint region. We also selected only voxels that produced a reliable response pattern to the stimuli across scanning sessions. Note that rather than using a simple correlation to determine reliability, we used the equation from Norman-Haignere et al. (2015) to measure the reliability across split halves of our data. This reliability measure differs from a Pearson correlation in that it assigns high values to voxels that respond consistently to sounds and does not penalize them even if their response does not vary much between sounds, which is the case for many voxels within primary auditory cortex:

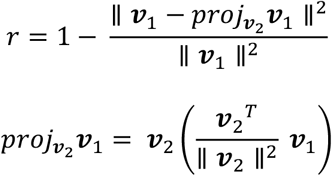

where ***ν***_1_ and ***ν***_2_ indicate the vector of beta weights from a single voxel for the 192 sounds, estimated separately for the two halves of the data (***ν***_1_ = first three repetitions from runs 1-24, ***ν***_2_ = last three repetitions from runs 25-48), and ∥ ∥ is the L2 norm. Note that these equations differ from Equations 1 and 2 in Norman-Haignere et al. (2015), because the equations as reported in the paper contained a typo: the L2-norm terms were not squared. We used the same reliability cutoff as in our prior study (r > 0.3). Of the sound-responsive voxels, 54.47% of them also met the reliability criteria. Using these two selection criteria, the median number of voxels per participant = 1,286, SD = 254 (**Supplemental Figure S1A**). The number of selected voxels did not differ significantly between musicians (median = 1,216, SD = 200) and non-musicians (median = 1,341, SD = 284; z = −1.40, p = 0.16, effect size r = −0.31, two-tailed Wilcoxon rank sum test), and the anatomical location of the selected voxels was largely similar across groups (**Supplemental Figure S1B**).

Unlike our prior study, we collected whole-brain data in this experiment and thus were able to repeat our analyses without any anatomical constraint. While a few additional voxels outside of the mask do meet our selection criteria (**Supplemental Figure S2**), the resulting components are very similar to those obtained using the anatomical mask, both in response profiles (**Supplemental Figure S3A**; correlations ranging from r = 0.91 to r > 0.99, SD = 0.03) and voxel weights (**Supplemental Figure S3B**).

### Decomposition algorithm

The decomposition algorithm approximates the response of each voxel (***ν***_*i*_) as the weighted sum of a small number of component response profiles that are shared across voxels (**Figure 1B**):

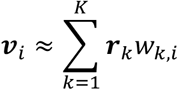

where ***r***_*k*_ represents the *k*^th^ component response profile that is shared across all voxels, *w*_*k*,*i*_ represents the weight of component *k* in voxel *i*, and *K* is the total number of components.

We concatenated the selected voxel responses from all participants into a single data matrix ***D*** (*S* sounds × *V* voxels). We then approximated the data matrix as the product of two smaller matrices: (1) a response matrix ***R*** (*S* sounds x *K* components) containing the response profile of all inferred components to the sound set, and (2) a weight matrix ***W*** (*K* components × *V* voxels) containing the contribution of each component response profile to each voxel. Using matrix notation this yields:

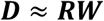

The method used to infer components was described in detail in our prior paper (Norman-Haignere et al., 2015) and code is available online (https://github.com/snormanhaignere/nonparametric-ica). The method is similar to standard algorithms for independent components analysis (ICA) in that it searches amongst the many possible solutions to the factorization problem for components that have a maximally non-Gaussian distribution of weights across voxels (the non-Gaussianity of the components inferred in this study can be seen in **Supplemental Figure S4**). The method differs from most standard ICA algorithms in that it maximizes non-Gaussianity by directly minimizing the entropy of the component weight distributions across voxels as measured by a histogram, which is feasible due to the large number of voxels. Entropy is a natural measure to minimize because the Gaussian distribution has maximal entropy. The algorithm achieves this goal in two steps. First, PCA is used to whiten and reduce the dimensionality of the data matrix. This was implemented using the singular value decomposition:

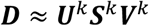

where ***U***^*k*^ contains the response profiles of the top *K* principal components (192 sounds × *K* components), ***V***^*k*^ contains the whitened voxel weights for these components (*K* components × 26,792 voxels), and ***S***^*k*^ is a diagonal matrix of singular values (*K* × *K*). The number of components (*K*) was chosen by measuring the amount of voxel response variance explained by different numbers of components and the accuracy of the components in predicting voxel responses in left-out data. Specifically, we chose a value of *K* that balanced these two measures such that the set of components explained a large fraction of the voxel response variance (which increases monotonically with additional components), but still maintained good prediction accuracy (which decreases once additional components begin to cause overfitting). In practice, the plateau in the amount of explained variance coincided with the peak of the prediction accuracy.

The principal component weight matrix is then rotated to maximize the negentropy (*J*) summed across components:

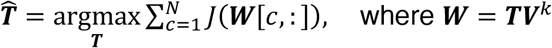

where ***W*** is the rotated weight matrix (*K* × 26,792), ***T*** is an orthonormal rotation matrix (*K* × *K*), and ***W***[*c*, : ] is the c^th^ row of ***W***. We estimated entropy using a histogram-based method (Moddemeijer, 1989) applied to the voxel weight vector for each component (***W***[*c*, : ]), and defined negentropy as the difference in entropy between the empirical weight distribution and a Gaussian distribution of the same mean and variance:

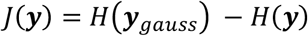

The optimization is performed by iteratively rotating pairs of components to maximize negentropy, which is a simple algorithm that does not require the computation of gradients, and is feasible for small numbers of components (the number of component pairs grows as 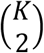).

This voxel decomposition analysis was carried out on three different data sets: i) on voxels from the 10 musicians only, ii) on voxels from the 10 non-musicians only, and iii) on voxels from all twenty participants. We note that the derivation of a set of components using this method is somewhat akin to a fixed-effects analysis, in that it concatenates participants’ data and infers a single set of components to explain the data from all participants at once. However, the majority of the analyses that we carried out using these components (as described below) involve deriving participant-specific metrics and investigating the consistency of effects across participants.

### Measuring component selectivity

To quantify the selectivity of the music component, we measured the difference in mean response profile magnitude between music and non-music sounds, divided by their pooled standard deviation (Cohen’s d). This measure provides a measure of the separability of the two sound categories within the response profile. We measured Cohen’s d for several different pairwise comparisons of sound categories. In each case, the significance of the separation of the two stimulus categories was determined using a permutation test (permuting stimulus labels between the two categories 10,000 times). This null distribution was then fit with a Gaussian, a p-value from which was assigned to the observed value of Cohen’s d.

### Anatomical component weight maps

To visualize the anatomical distribution of component weights, individual participants’ component weights were projected onto the cortical surface of the standard Freesurfer FsAverage template, and a random effects analysis (t-test) was performed to determine whether component weights were significantly greater than zero across participants at each voxel. To visualize the component weight maps separately for musicians and non-musicians, a separate random effects analysis was run for the participants in each group. To correct for multiple comparisons, we adjusted the false discovery rate (FDR, c(V) = 1, q = 0.05) using the method from Genovese, Lazar, and Nichols (Genovese et al., 2002).

We note that the details of the analyses and plotting conventions used for visualizing component weight maps differ from those of our previous study (Norman-Haignere et al., 2015). These differences include the process involved in aggregating weights across subjects (Norman-Haignere et al. smoothed individual participants’ data and then averaged across participants, whereas there was no smoothing or averaging across participants in the current study), the use of different measures of statistical significance of the component weights (a random effects analysis across subjects in the current study, a permutation test across sounds in Norman-Haignere et al.), different thresholding (an FDR threshold of q = 0.05 in the current study, whereas the component weight maps in Norman-Haignere et al. showed the entire weight distribution and were not thresholded). We made these changes in the current study so that we could ask about the consistency of effects across participants, and better visualize which voxels’ component weights passed standard significance thresholds. However, we note that these changes cause the maps we regenerated from the Norman-Haignere et al. (2015) data (**Figure 3C** & **D**) to look somewhat different from those shown in the original paper (**Figure 5A**).

### Component voxel weights within anatomical ROIs

In addition to projecting the weight distributions on the cortical surface, we summarized their anatomical distribution by measuring the mean component voxel weight within a set of standardized anatomical ROIs. To create these ROIs, a set of fifteen parcels were selected from an atlas (Glasser et al.2016) to fully encompass the superior temporal plane and superior temporal gyrus (STG). To identify a small set of ROIs suitable for evaluating the music component weights in our current study, we superimposed these anatomical parcels onto the weights of the music component from our previously published study (Norman-Haignere et al., 2015; shown in **Figure 5A**), and then defined ROIs by selecting sets of the anatomically-defined parcels that best correspond to regions of high vs. low music component weights (**Figure 5B**). The mean component weights within these ROIs were computed separately for each participant, and then averaged across participants for visualization purposes (e.g. **Figure 5C**). We then ran a 4 (ROI) x 2 (hemisphere) repeated-measures ANOVA on these weights. A separate ANOVA was run for musicians and non-musicians, to evaluate each group separately.

To compare the magnitude of the main effect of ROI with the main effect of hemisphere, we bootstrapped across participants, resampling participants 1,000 times. We re-ran the repeated-measures ANOVA on each sample, each time measuring the difference in the effect size for the two main effects, i.e. η_pROI_^2^ - η_pHemi_^2^. We then calculated the 95% CI of this distribution of effect size differences. The significance of the difference in main effects was evaluated by determining whether or not each group’s 95% CI for the difference overlapped with zero.

In addition, we ran a Bayesian repeated-measures ANOVA on these same data, implemented in JASP v.0.13.1, using the default prior (Cauchy distribution, r = 0.5). Effects are reported as the Bayes Factor for inclusion (BF_inc_) of each main effect and/or interaction, which is the ratio between the likelihood of the data given the model including the effect in question vs. the likelihood of the next simpler model without the effect in question.

## RESULTS

Our primary question was whether cortical music selectivity is present in people with almost no musical training. To that end, we scanned ten people with almost no musical training (**Table 1**), and used voxel decomposition to isolate music-selective neural populations. We also included another set of ten participants with extensive musical training for comparison. First, we asked whether the response components across all of auditory cortex reported by Norman-Haignere et al. (2015) replicate in both non-musicians and highly trained musicians when analyzed separately. Second, we examined the detailed characteristics of music selectivity in particular, to test whether its previously documented key properties are present in both non-musicians and highly trained musicians. Third, we took advantage of our expanded stimulus set to look at additional properties of music selectivity, such as the response to musical genres that are less familiar to our Western participants.

### Replication of functional components of auditory cortex from Norman-Haignere et al. (2015) in musicians and in non-musicians

#### Replication of previous voxel decomposition results

We first tested the extent to which we would replicate the overall functional organization of auditory cortex reported by Norman-Haignere et al. (2015) in people with almost no musical training, using the voxel decomposition method introduced in that paper. We also performed the same analysis on a group of highly trained musicians. Specifically, in every participant, we measured the response of voxels within auditory cortex to 192 natural sounds (**Figure 1A** & **B;** the average response of each voxel to each sound was estimated using a standard hemodynamic response function). Then, separately for non-musicians and musicians, we used voxel decomposition to model the response of these voxels as the weighted sum of a small number of canonical response components (**Figure 1C**). This method factorizes the voxel responses into two matrices: one containing the components’ response profiles across the sound set, and the second containing voxel weights specifying the extent to which each component contributes to the response of each voxel.

Since the only free parameter in this analysis is the number of components recovered, the optimal number of components was determined by measuring the fraction of the reliable response variance the components explain. In the previous study (Norman-Haignere et al., 2015), six components were sufficient to explain over 80% of the reliable variance in voxel responses. We found the same to be true in both participant groups of the current study: six components were needed to optimally model the data from the 10 participants in separate analyses of each group. The six components explained 88.56% and 88.09% of the reliable voxel response variance for non-musicians and musicians respectively, after which the amount of explained variance for each additional component plateaued (**Supplemental Figure S5**).

Next, we examined the similarity of the components inferred from non-musicians to the components from our previous study, comparing their responses to the 165 sounds common to both. Because the order of the components inferred using ICA holds no significance, we first used the Hungarian algorithm (Kuhn, 1955) to optimally reorder the components, maximizing their correlation with the components from our previous study. For comparisons of the response profile matrices of two groups of subjects, we matched components using the weight matrices; conversely, for comparisons involving the voxel weights, we matched components using the response profile matrices (see Methods). In practice, the component matches were identical regardless of which matrices were used for matching. We also conducted the same analysis for the musicians, comparing the components derived from their data to those in our previous study.

For both non-musicians and musicians, corresponding pairs of components were highly correlated with those from our previous study, with r-values for non-musicians’ components ranging from 0.63 to 0.99 (**Figure 2A** & **B**) and from 0.66 to 0.99 for musicians’ components (**Figure 2C** & **D**).

**Figure 2.**
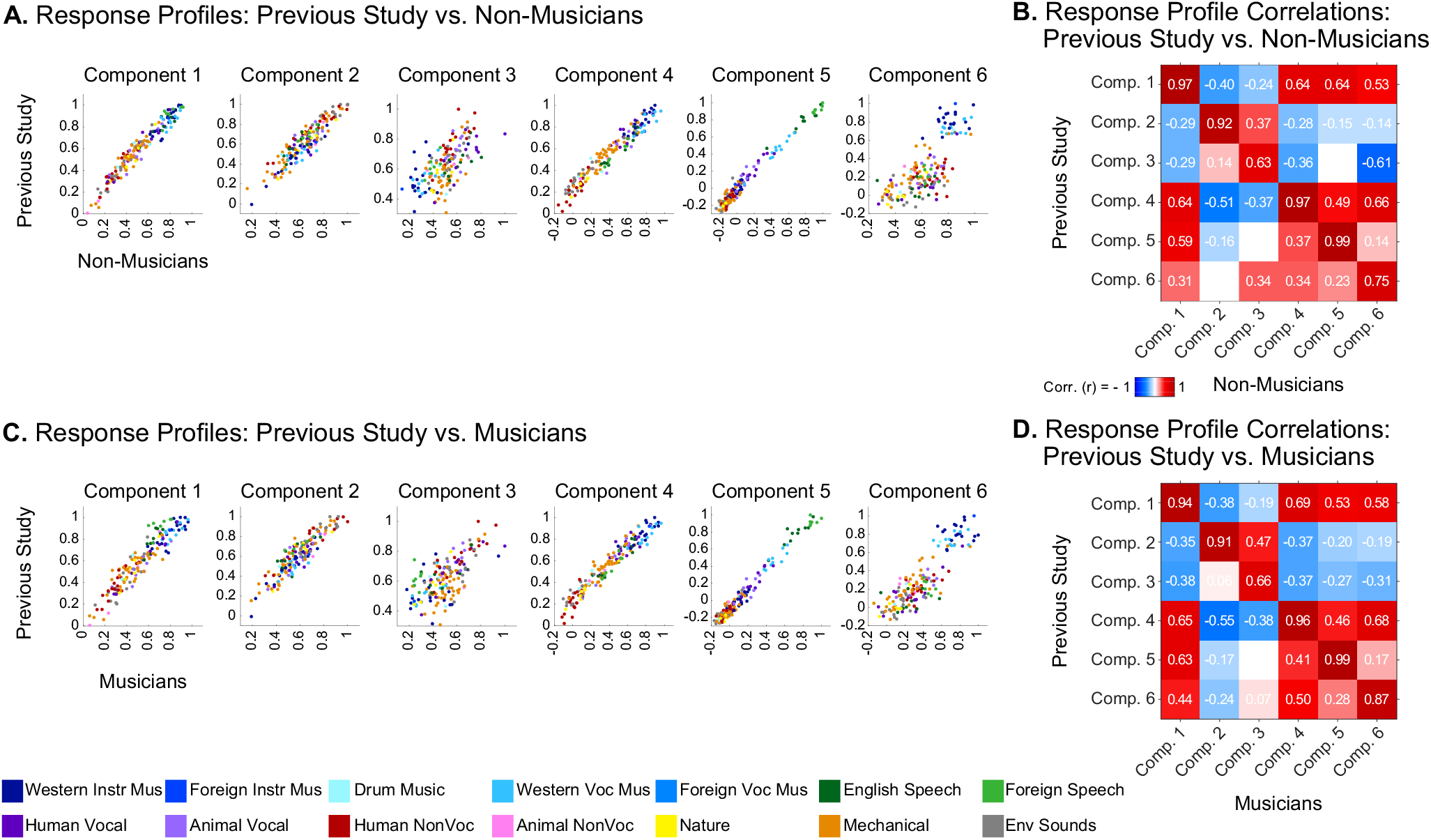
Replication of components from Norman-Haignere et al. (2015). (**A**) Scatterplots showing the correspondence between the component response profiles from the previous study (y-axis) and those inferred from non-musicians (x-axis). Sounds are colored according to their semantic category, as determined by raters on Amazon Mechanical Turk. Note that the axes differ slightly between groups in order to make it possible to clearly compare the pattern of responses across sounds independent of the overall response magnitude. (**B**) Correlation matrix comparing component response profiles from the previous study (y-axis) and those inferred from non-musicians (x-axis). (**C, D**) Same as **A & B** but for musicians.

#### Component response profiles and selectivity for sound categories

Four of the six components from our previous study captured expected acoustic properties of the sound set (e.g. frequency, spectrotemporal modulation; see **Supplemental Figure S6A** for analyses relating the responses of these components to audio frequency and spectrotemporal modulation) and were concentrated in and around primary auditory cortex (PAC), consistent with prior results (Chi et al., 2005; Schönwiesner and Zatorre, 2009; Humphries et al., 2010; Da Costa et al., 2011; Herdener et al., 2013; Santoro et al., 2014; Hullett et al., 2016; Norman-Haignere and McDermott, 2018). The two remaining components responded selectively to speech (**Figure 3A**, left column) and music (**Figure 3B**, left column), respectively, and were not well accounted for using acoustic properties alone (**Supplemental Figure S6A**). The corresponding components inferred from non-musicians (**Figure 3A** & **B**, middle columns) and musicians (**Figure 3A** & **B**, right columns) also show this category selectivity.

**Figure 3.**
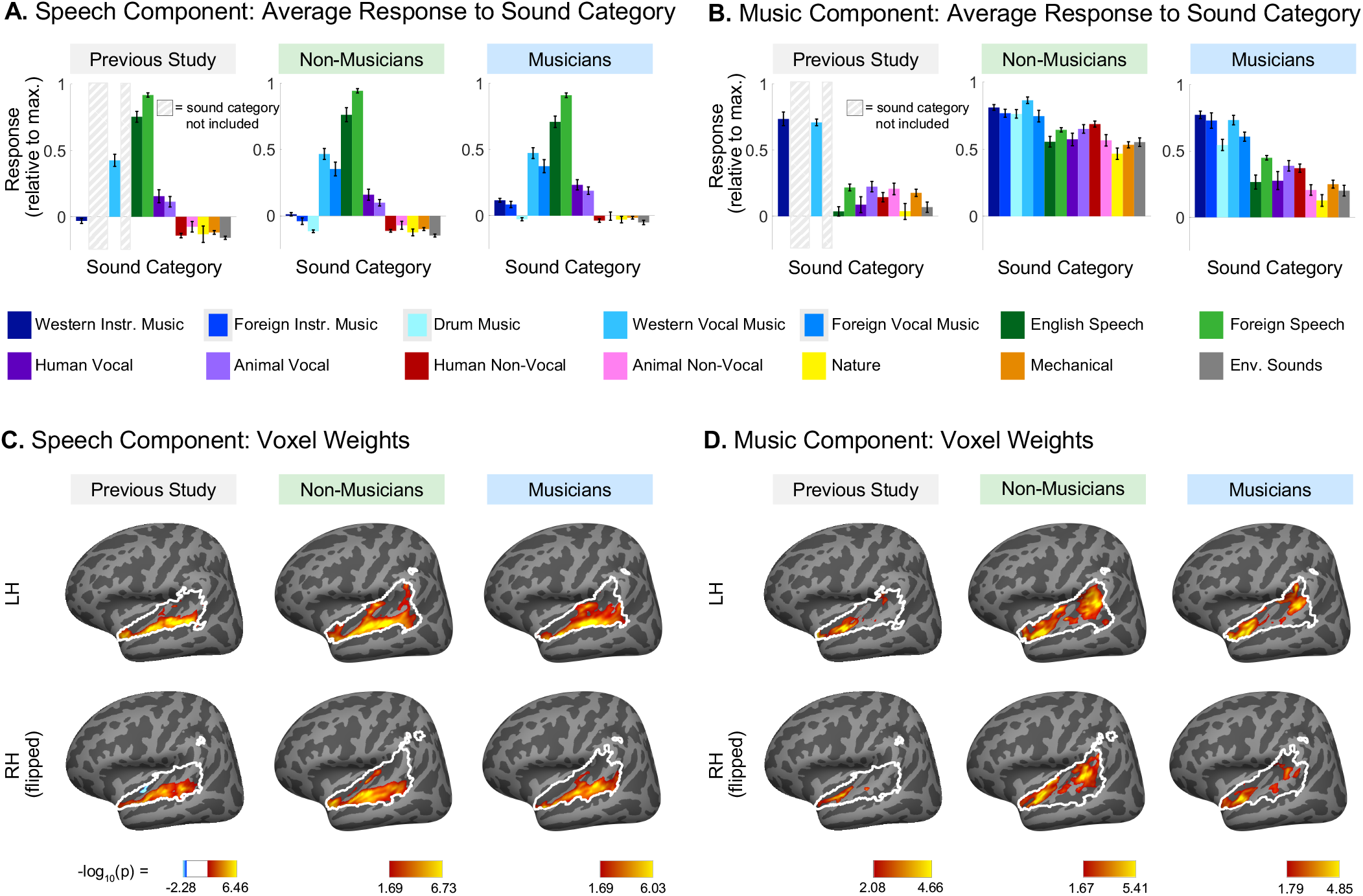
Comparison of speech-selective and music-selective components for participants from previous study, non-musicians, and musicians. Component response profiles averaged by sound category (as determined by raters on Amazon Mechanical Turk). (**A**) The speech-selective component responds highly to speech and music with vocals, and minimally to all other sound categories. Shown separately for the previous study (left), non-musicians (middle) and musicians (right). Note that the previous study contained only a subset of the stimuli used in the current study, so some conditions were not included and are thus replaced by a gray rectangle in the plots and surrounded by a gray rectangle in the legend. (**B**) The music-selective component (right) responds highly to both instrumental and vocal music, and less strongly to other sound categories. Note that “Western Vocal Music” stimuli were sung in English. We note that the mean response profile magnitude differs between groups, but that selectivity as measured by separability of music and non-music is not affected by this difference (see text for explanation). For both **A** and **B**, error bars plot one standard error of the mean across sounds from a category, computed using bootstrapping (10,000 samples). (**C**) Spatial distribution of speech-selective component voxel weights in both hemispheres. (**D**) Spatial distribution of music-selective component voxel weights. Color denotes the statistical significance of the weights, computed using a random effects analysis across subjects comparing weights against 0; p-values are logarithmically transformed (−log_10_[p]). The white outline indicates the voxels that were both sound-responsive (sound vs. silence, p < 0.001 uncorrected) and split-half reliable (r > 0.3) at the group level (see Methods for details). The color scale represents voxels that are significant at FDR q = 0.05, with this threshold computed for each component separately. Voxels that do not survive FDR correction are not colored, and these values appear as white on the color bar. The right hemisphere (bottom rows) is flipped to make it easier to visually compare weight distributions across hemispheres. Note that the secondary posterior cluster of music component weights is not as prominent in this visualization of the data from Norman-Haignere et al. (2015) due to the thresholding procedure used here, and we found in additional analyses that a posterior cluster emerged if a more lenient threshold is used.

We note that the mean response profile magnitude for the music-selective component differed between groups, being lower in musicians than non-musicians (**Figure 3B**). This effect seems unlikely to be a consequence of musical training because the mean response magnitude was lower still in the previous study, whose participants had substantially less musical training than the musicians in the current study. Further, we have found that component response profile magnitude tends to vary depending on the method used to infer the components. For example, using a probabilistic parametric matrix factorization model instead of the simpler, non-parametric method presented throughout this paper resulted in components that had different mean responses despite otherwise being very similar to those obtained via ICA (see Supplemental Information for details of the parametric model, and **Supplemental Figure S7** for the component response profiles inferred using this method). Moreover, the information about music contained in the response as measured by the separability of music versus non-music sounds (Cohen’s d) is independent of this overall mean response (see **Figure 4**). For these reasons we do not read much into this apparent difference between groups.

**Figure 4.**
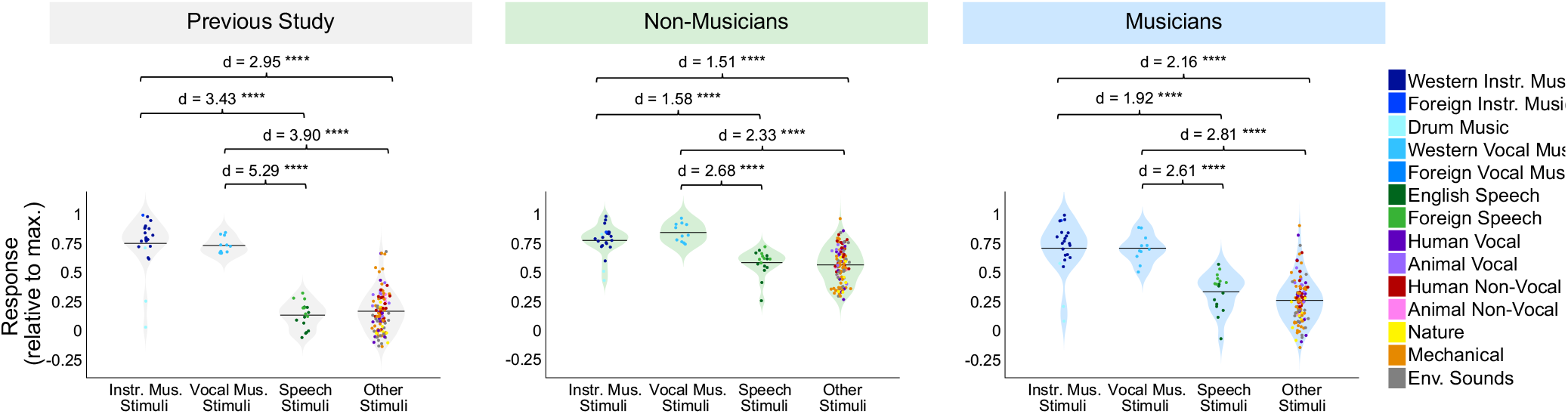
Separability of sound categories in music-selective components of non-musicians and musicians. Distributions of (1) instrumental music stimuli, (2) vocal music stimuli, (3) speech stimuli, and (4) other stimuli within the music component response profiles from our previous study (left, grey shading), as well as those inferred from musicians (center, blue shading) and non-musicians (right, green shading). The mean for each stimulus category is indicated by the horizontal black line. The separability between pairs of stimulus categories (as measured using Cohen’s d) is shown above each plot. Individual sounds are colored according to their semantic category. See **Table 2** for results of pairwise comparisons indicated by brackets; **** = significant at p < 0.0001 two-tailed.

We also found similarities in the anatomical distribution of speech- and music-selective component weights between the previous study and both groups in the current study. The weights for the speech-selective component were concentrated in the middle portion of the superior temporal gyrus (midSTG, **Figure 3C**), as expected based on previous reports (Scott et al., 2000; Hickok and Poeppel, 2007; Overath et al., 2015). In contrast, the weights for the music-selective component were most prominent anterior to PAC in the planum polare, with a secondary cluster posterior to PAC in the planum temporale (**Figure 3D**) (Ohnishi et al., 2001; Margulis et al., 2009; Dick et al., 2011; Fedorenko et al., 2012; Angulo-Perkins et al., 2014; Armony et al., 2015; Norman-Haignere et al., 2015).

Together, these findings show that we are able to twice replicate the overall component structure underlying of auditory cortical responses described in our previous study (once for non-musicians and once for musicians). Further, both non-musicians and musicians show category-selective components that are largely similar to those in our previous study, including a single component that appears to be selective for music.

### Characterizing music selectivity in non-musicians and musicians separately

We next asked whether nonmusicians exhibited the signature response characteristics of music selectivity documented in our prior paper: (1) the music component response profile showed a high response to both instrumental and vocal music and a low response to all other categories, including speech, (2) music component voxel weights were highest in anterior superior temporal gyrus (STG), with indications of a secondary concentration of weights in posterior STG, and low weights in both PAC and lateral STG, and (3) music component voxel weights had a largely bilateral distribution. We examined whether each of these properties was present in the components separately inferred from non-musicians and musicians.

#### Response profiles show selectivity for both instrumental and vocal music

The defining feature of the music-selective component from Norman-Haignere et al. (2015) was that its response profile showed a very high response to stimuli that had been categorized by humans as “music”, including both instrumental music and music with vocals, relative to non-music stimuli, including speech (**Figure 3B**, left column). The category-averaged responses of the music-selective component showed similar effects in non-musicians and highly trained musicians (**Figure 3B**, center and right columns).

To quantify this music selectivity, we measured the difference in mean response profile magnitude between music and non-music sounds, divided by their pooled standard deviation (Cohen’s d). So that we could compare across our previous and current experiments, this was done using the set of 165 sounds that were common to both studies. We measured Cohen’s d separately for several different pairwise comparisons: (1) instrumental music vs. speech stimuli, (2) instrumental music vs. other non-music stimuli, (3) vocal music vs. speech stimuli, and (4) vocal music vs. other non-music stimuli (**Figure 4**). In each case, the significance of the separation of the two stimulus categories was determined using a nonparametric test permuting stimulus labels 10,000 times. All four of these statistical comparisons were highly significant for non-musicians when analyzed separately (all p’s < 10^−5^, **Table 2**). This result shows that the music component is highly music selective in non-musicians, in that it responds highly to both instrumental and vocal music, and significantly less to both speech and other non-music sounds. Similar results were also found for musicians (all p’s < 10^−5^, **Table 2**). We note that the selectivity of the music component inferred from non-musicians seems to be slightly lower than that of the component inferred from musicians, but we are not sufficiently powered to directly test for differences in selectivity between groups (see “Direct group comparisons of music selectivity” in the **Supplemental Information**). It’s also true that the values of Cohen’s d tend to be somewhat larger for the music component from Norman-Haignere et al. (2015) than for the components inferred from both groups of participants in the current study. It is not clear why this is the case, but it is more likely to be due to slight methodological differences between the experiments than an effect of musical training, because the participants in our previous study had intermediate levels of musical training.

**Table 2.** Results of pairwise comparisons between stimulus categories shown in **Figure 4B**. The significance of the separation of the two stimulus categories was determined using a nonparametric test permuting stimulus labels 10,000 times.

|  |  | Stimulus Category | Mean | SD | Cohen's d | p-value |
| --- | --- | --- | --- | --- | --- | --- |
| NON-MUSICIANS | 1 | Instrumental Music | 0.78 | 0.13 | 1.58 | 2.14E-06 |
|  |  | Speech | 0.59 | 0.11 |  |  |
|  | 2 | Instrumental Music | 0.78 | 0.13 | 1.51 | 1.88E-10 |
|  |  | Other | 0.57 | 0.15 |  |  |
|  | 3 | Vocal Music | 0.85 | 0.08 | 2.68 | 4.54E-11 |
|  |  | Speech | 0.59 | 0.11 |  |  |
|  | 4 | Vocal Music | 0.85 | 0.08 | 2.33 | 1.87E-12 |
|  |  | Other | 0.57 | 0.15 |  |  |
| MUSICIANS | 1 | Instrumental Music | 0.72 | 0.22 | 1.92 | 1.19E-08 |
|  |  | Speech | 0.34 | 0.16 |  |  |
|  | 2 | Instrumental Music | 0.72 | 0.22 | 2.16 | 7.40E-20 |
|  |  | Other | 0.27 | 0.19 |  |  |
|  | 3 | Vocal Music | 0.72 | 0.12 | 2.61 | 8.38E-11 |
|  |  | Speech | 0.34 | 0.16 |  |  |
|  | 4 | Vocal Music | 0.72 | 0.12 | 2.81 | 3.62E-18 |
|  |  | Other | 0.27 | 0.19 |  |  |

#### Music component weights concentrated in anterior and posterior STG

A second notable property of the music component from Norman-Haignere et al. (2015) was that the weights were concentrated in distinct regions of non-primary auditory cortex, with the most prominent cluster located in anterior STG, and a secondary cluster located in posterior STG (at least in the left hemisphere). Conversely, music component weights were low in primary auditory cortex (PAC) and intermediate in non-primary lateral STG (see **Figure 5A**).

**Figure 5.**
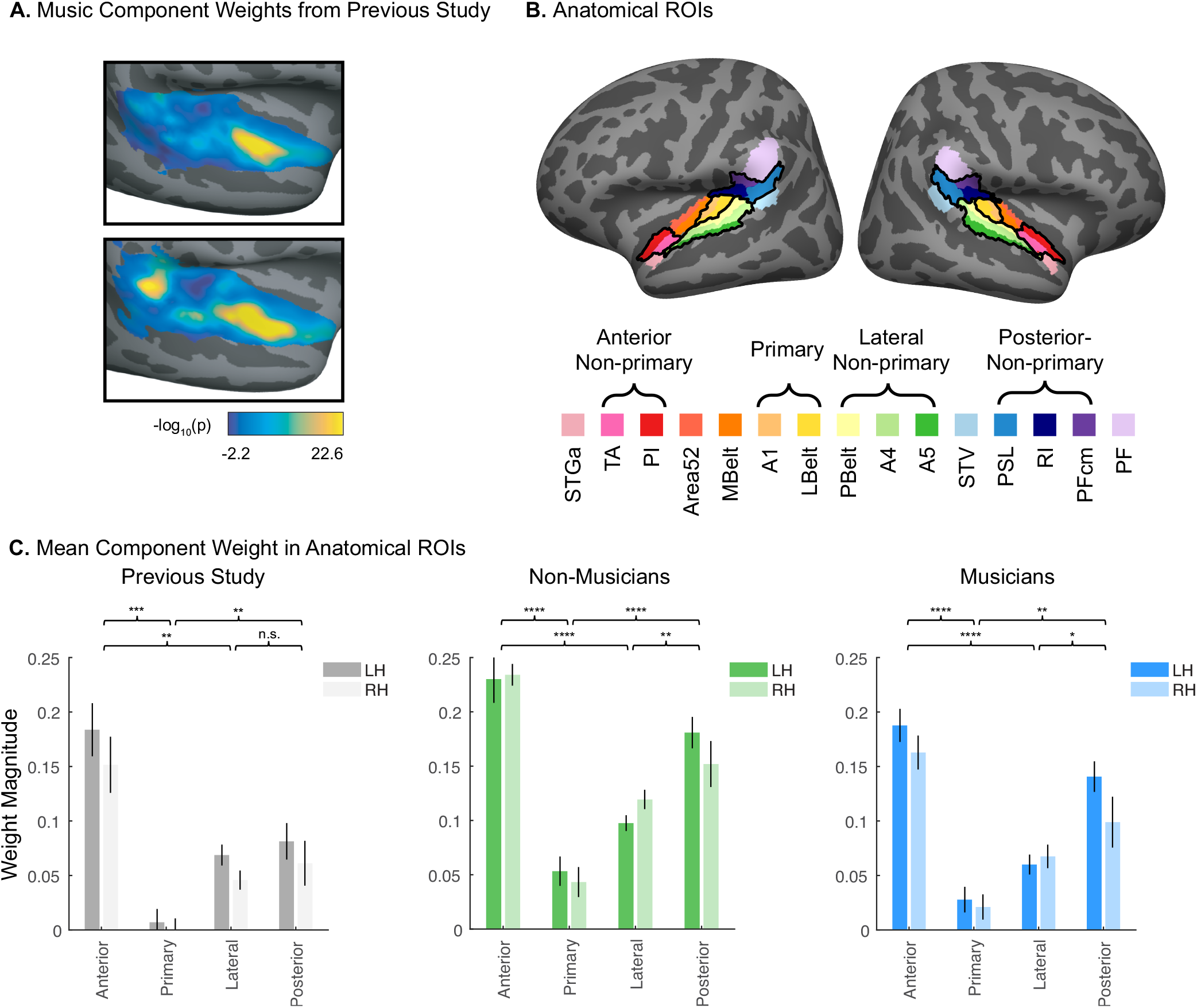
Quantification of bilateral anterior/posterior concentration of voxel weights for the music-selective components inferred in non-musicians and musicians separately. (A) Music component voxel weights, reproduced from Norman-Haignere et al. (2015). See Methods for details concerning the analysis and plotting conventions from our previous paper. (B) Fifteen standardized anatomical parcels were selected from Glasser et al. (2016), chosen to fully encompass the superior temporal plane and superior temporal gyrus (STG). To come up with a small set of ROIs to use to evaluate the music component weights in our current study, we superimposed these anatomical parcels onto the weights of the music component from our previously published study (Norman-Haignere et al., 2015), and then defined ROIs by selecting sets of the anatomically-defined parcels that correspond to regions of high (anterior non-primary, posterior non-primary) vs. low (primary, lateral non-primary) music component weights. The anatomical parcels that comprise these four ROIs are indicated by the brackets, and outlined in black on the cortical surface. (C) Mean music component weight across all voxels in each of the four anatomical ROIs, separately for each hemisphere, and separately for non-musicians (left, green shading) and musicians (right, blue shading) and A repeated-measures ROI x hemisphere ANOVA was conducted for each group separately. Error bars plot one standard error of the mean across participants. Brackets represent pairwise comparisons that were conducted between ROIs with expected high vs. low component weights, averaged over hemisphere. See **Table 3** for full results of pairwise comparisons, and **Supplemental Figure S8** for component weights from all fifteen anatomical parcels; * = significant at p < 0.05 two-tailed, ** = significant at p < 0.01 two-tailed, *** = significant at p < 0.001 two-tailed, **** = significant at p < 0.0001 two-tailed. Note that because of our prior hypotheses and the significance of the omnibus F-test, we did not correct for multiple comparisons.

To assess whether these anatomical characteristics were evident for the music components inferred from our non-musician and musician participants, we superimposed standardized anatomical parcels (Glasser et al., 2016) on the data, and defined anatomical regions of interest (ROIs) by selecting four sets of these anatomically-defined parcels that best corresponded to anterior non-primary, primary, and lateral non-primary auditory cortex (**Figure 5B**). We then calculated the average music component weight for each individual participant within each of these four anatomical ROIs, separately for each hemisphere (**Figure 5C**). This was done separately for non-musicians and musicians, using their respective music components. We did not perform this analysis on the data from our previous study because its anatomical coverage was more restricted than the data we collected here.

For each group, a 4 (ROI) x 2 (hemisphere) repeated measures ANOVA on these mean component weights showed a significant main effect of ROI for both non-musicians (F(3, 27) = 50.12, p = 3.63e-11, η_p_^2^ = 0.85) and musicians (F(3, 27) = 19.62, p = 5.90e-07, η_p_^2^ = 0.69). Pairwise comparisons showed that for each group, component weights were significantly higher in the anterior and posterior non-primary ROIs than both the primary and lateral non-primary ROIs when averaging over hemispheres (**Table 3**; non-musicians: all p’s < 0.004, musicians: all p’s < 0.03).

**Table 3.** Results of pairwise comparisons between mean weights in ROIs shown in **Figure 5C**. Component weights were first averaged over hemispheres, and significance between ROI pairs was evaluated using paired t-tests. Note that because of our prior hypotheses and the significance of the omnibus F-test,, we did not correct for multiple comparisons.

|  |  | ROI | t-Value | Cohen's d | p-value |
| --- | --- | --- | --- | --- | --- |
| NON-MUSICIANS | 1 | Anterior non-primary<br>Primary | 15.91 | 5.03 | 6.75E-08 |
|  | 2 | Anterior non-primary<br>Lateral non-primary | 8.99 | 2.84 | 8.61E-06 |
|  | 3 | Posterior non-primary<br>Primary | 7.25 | 2.29 | 4.81E-05 |
|  | 4 | Posterior non-primary<br>Lateral non-primary | 3.84 | 1.21 | 0.0040 |
| MUSICIANS | 1 | Anterior non-primary<br>Primary | 11.40 | 3.61 | 1.19E-06 |
|  | 2 | Anterior non-primary<br>Lateral non-primary | 7.66 | 2.42 | 3.14E-05 |
|  | 3 | Posterior non-primary<br>Primary | 4.49 | 1.42 | 0.0015 |
|  | 4 | Posterior non-primary<br>Lateral non-primary | 2.72 | 0.86 | 0.0237 |

These results show that in both non-musicians and musicians, music selectivity is concentrated in anterior and posterior STG, and present to a lesser degree in lateral STG, and only minimally in PAC.

#### Music component weights are bilaterally distributed

A third characteristic of the previously described music selectivity is that it was similarly distributed across hemispheres, with no obvious lateralization (Norman-Haignere et al., 2015). The repeated-measures ANOVA described above showed no evidence of lateralization in either non-musicians or musicians (**Figure 5C**; non-musicians: F(1, 9) = 0.15, p = 0.71, η_p_^2^ = 0.02; musicians: F(1, 9) = 2.43, p = 0.15, η_p_^2^ = 0.21). Furthermore, for both groups, the effect size of ROI within a hemisphere was significantly larger than the effect size of hemisphere (measured by bootstrapping across participants to get 95% CIs around the difference in the effect size for the two main effects, i.e. η_pROI_^2^ – η_pHemi_^2^; the significance of the difference in main effects was evaluated by determining whether or not each group’s 95% CI for the difference overlapped with zero: non-musicians’ CI = [0.37 0.89], musicians’ CI = [0.16 0.82]).

Because the lack of a significant main effect of hemisphere could be due to insufficient statistical power, we ran a Bayesian version of the repeated-measures ANOVA, which allows us to quantify evidence both for and against the null hypothesis that there was not a main effect of hemisphere. We used JASP (JASP Team, 2020), with its default prior (Cauchy distribution, r = 0.5), and computed the Bayes Factor for inclusion of each main effect and/or interaction (the ratio between the likelihood of the data given the model including the effect in question vs. the likelihood of the next simpler model without the effect in question, with values further from 1 providing stronger evidence in favor of one model or the other). We found no evidence for a main effect of hemisphere (Bayes factor of inclusion, BF_incl_ = 0.77 for musicians, suggestive of anecdotal evidence against inclusion; BF_incl_ = 0.24 for non-musicians, suggestive of moderate evidence against inclusion, using the guidelines suggested by Lee & Wagenmakers, 2013). By contrast, the main effect of ROI was well supported (BF_incl_ for ROI for non-musicians = 1.02e17, and for musicians = 6.11e12, both suggestive of extreme evidence in favor of inclusion).

Neither group showed a significant ROI x hemisphere interaction (non-musicians: F(3, 27) = 1.48, p = 0.24, η_p_^2^ = 0.14; musicians: F(3, 27) = 2.24, p = 0.11, η_p_^2^ = 0.20). This was also the case for the Bayesian repeated-measures ANOVA, in which the Bayes Factors for the interaction between ROI and hemisphere provided anecdotal evidence against including the interaction term in the models (BF_incl_ = 0.38 for non-musicians, BF_incl_ = 0.36 for musicians).

The fact that the music component inferred from non-musicians exhibits all of the previously described features of music selectivity (Norman-Haignere et al., 2015) suggests that explicit musical training is not necessary for a music-selective neural population to arise in the human brain. In addition, the results from musicians suggest that that the signature properties of music selectivity are not drastically altered by extensive musical experience. Both groups exhibited a single response component selective for music. And in both groups, this selectivity was present for both instrumental and vocal music, was localized to anterior and posterior non-primary auditory cortex, and was present bilaterally.

### New insights into music selectivity: Music-selective regions of auditory cortex show high responses to drum rhythms and unfamiliar musical genres

Because our experiment utilized a broader stimulus set than the original study (Norman-Haignere et al., 2015), we were able to use the inferred components to ask additional questions about the effect of experience on music selectivity, as well as gain new insights into the nature of cortical music selectivity. The set of natural sounds used in this study included a total of 62 music stimuli, spanning a variety of instruments, genres, and cultures. Using this diverse set of music stimuli, we can begin to address the questions of (1) whether music selectivity is specific to the music of one’s own culture, and (2) whether music selectivity is driven solely by features related to pitch, like the presence of a melody. Here we analyze the music component inferred from all 20 participants since similar music components were inferred separately from musicians and non-musicians (see **Supplemental Figure S9** for details of the components inferred from all 20 participants).

To expand beyond the original stimulus set from Norman-Haignere et al. (2015), which contained music exclusively from traditionally Western genres and artists, we selected additional music clips from several non-Western musical cultures that varied in tonality and rhythmic complexity (e.g. Indian raga, Balinese gamelan, Chinese opera, Mongolian throat singing, Jewish klezmer, Ugandan lamellophone music) (**Figure 6A**). We expected that our American participants would have less exposure to these musical genres, allowing us to see whether the music component makes a distinction between familiar and less familiar music. The non-Western music stimuli were rated by American Mechanical Turk participants as being similarly musical (mean rating on 1-100 scale for Western music = 86.28, SD = 7.06; Non-Western music mean = 79.63, SD = 9.01; p = 0.37, 10,000 permutations) but less familiar (mean rating on 1-100 scale for Western music = 66.50, SD = 8.23; Non-Western music mean = 45.50, SD = 15.83; p < 1.0e-5, 10,000 permutations) than typical Western music. Despite this difference in familiarity, the magnitude of non-Western music stimuli within the music component was only slightly smaller than the magnitude of Western music stimuli (Cohen’s d = 0.64), a difference that was only marginally significant (**Figure 6B**; p = 0.052, nonparametric test permuting music stimulus labels 10,000 times). Moreover, the magnitudes of both Western and non-Western music stimuli were both much higher than non-music stimuli (Western music stimuli vs. non-music stimuli: Cohen’s d = 3.79, p < 0.0001, 10,000 permutations; non-Western music vs. non-music: Cohen’s d = 2.85; p < 0.0001, 10,000 permutations). Taken together, these results suggest that music-selective responses in auditory cortex occur even for relatively unfamiliar musical systems and genres.

**Figure 6.**
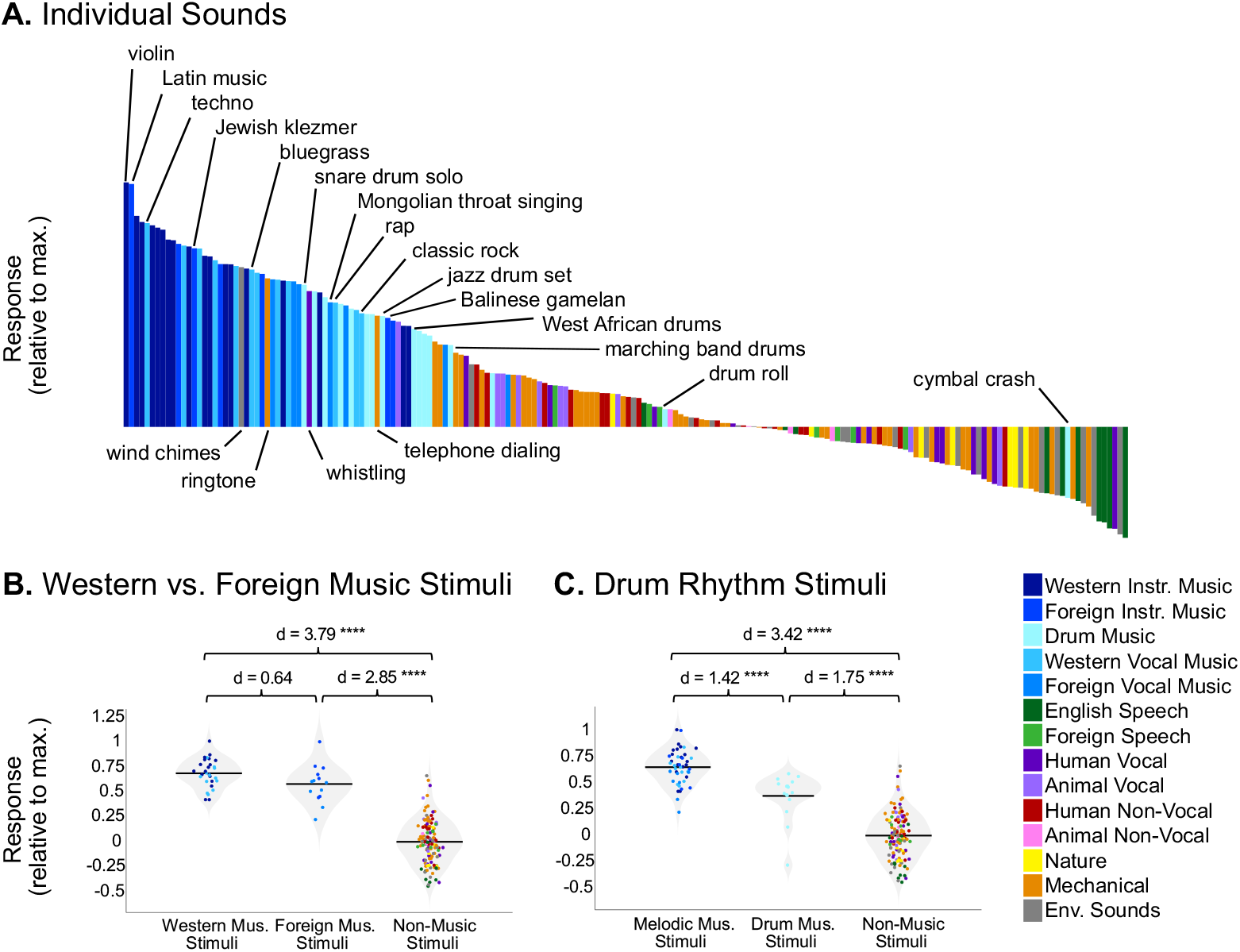
(**A**) Close-up of the response profile (192 sounds) for the music component inferred from all participants (n = 20), with example stimuli labeled. Note that there are a few “non-music” stimuli (categorized as such by Amazon Mechanical Turk raters) with high component rankings, but that these are all arguably musical in nature (e.g. wind chimes, ringtone). Conversely, “music” stimuli with low component rankings (e.g. “drumroll” and “cymbal crash”) do not contain salient melody or rhythm, despite being classified as “music” by human listeners. (**B**) Distributions of Western music stimuli (left), non-Western music stimuli (middle), and non-music stimuli (right) within the music component response profile inferred from all 20 participants, with the mean for each stimulus group indicated by the horizontal black line. The separability between categories of stimuli (as measured using Cohen’s d) is shown above the plot. Note that drum stimuli were left out of this analysis. (**C**) Distributions of melodic music stimuli (left), drum rhythm stimuli (middle), and non-music stimuli (right) within the music component response profile inferred from all 20 participants, with the mean for each stimulus group indicated by the horizontal black line. The separability between categories of stimuli (as measured using Cohen’s d) is shown above the plot, and significance was evaluated using a nonparametric test permuting stimulus labels 10,000 times; **** = significant at p < 0.0001 two-tailed. Sounds are colored according to their semantic category.

Which stimulus features drive music selectivity? One of the most obvious distinctions is between melody and rhythm. While music typically involves both melody and rhythm, when assembling our music stimuli we made an attempt to pick clips that varied in the prominence and complexity of their melodic and rhythmic content. In particular, we included 13 stimuli consisting of drumming from a variety of genres and cultures, because drum music mostly isolates the rhythmic features of music while minimizing (though not eliminating) melodic features. Whether music-selective auditory cortex would respond highly to these drum stimuli was largely unknown, partially because the Norman-Haignere et al. (2015) study only included two drum stimuli, one of which was just a stationary snare drum roll that produced a low response in the music component, likely because it lacks both musical rhythm and pitch structure. However, the drum stimuli in our study ranked relatively high in the music component response profile, averaging below the other instrumental and vocal music category responses (Cohen’s d = 1.42, p < 8.76e-07), but higher than the other non-music stimulus categories (Cohen’s d = 1.75, p < 9.60e-11) (**Figure 6C**). This finding suggests that the music component is not simply tuned to melodic information, but is also sensitive to rhythm.

## DISCUSSION

Our results show that cortical music selectivity is present in non-musicians and hence does not require explicit musical training to develop. Indeed, the same six response components that characterized human auditory cortical responses to natural sounds in our previous study were replicated twice here, once in non-musicians, and once in musicians. Our goal in this study was not to make statistical comparisons between non-musicians and musicians (which would have required a prohibitive amount of data, see “Direct group comparisons of music selectivity” in the **Supplemental Information**) but rather to assess whether the key properties of music selectivity were present in each group. Thus, while we cannot rule out the possibility that there are some differences between music-selective neural responses in musicians and non-musicians, we have shown that in both groups, voxel decomposition produced a single music-selective component, which was selective for both instrumental and vocal music, and which was concentrated bilaterally in anterior and posterior superior temporal gyrus (STG). We also observed that the music-selective component responds strongly to both drums and less familiar non-Western music. Together, these results show that passive exposure to music is sufficient for the development of music selectivity in non-primary auditory cortex, and that music-selective responses extend to rhythms with little melody, and to relatively unfamiliar musical genres.

### Origins of music selectivity

Our finding of music-selective responses in non-musicians is inconsistent with the hypothesis that explicit training is necessary for the emergence of music selectivity in auditory cortex, and suggests rather that music selectivity is either present from birth or results from passive exposure to music. If present from birth, music selectivity could in principle represent an evolutionary adaptation for music, definitive evidence for which has long been elusive (Darwin, 1871). But it is also plausible that music-specific representations emerge over development due to the behavioral importance of music in everyday life. For example, optimizing a neural network model to solve ecological speech and music tasks yields separate processing streams for the two tasks (Kell et al., 2018), suggesting that musical tasks sometimes require music-specific features. Another possibility is that music-specific features might emerge in humans or machines without tasks per se, due to the fact that music is acoustically distinct from other natural sounds. One way of testing this hypothesis might be to use generic unsupervised learning, for instance for producing efficient representations of sound (Lewicki, 2002; Carlson et al., 2012; Młynarski and McDermott, 2019), which might produce a set of features that are activated primarily by musical sounds.

Nearly all of our participants reported listening to music on a daily basis, and in other contexts this everyday musical experience has clear effects (Bigand, 1983; Bigand and Pineau, 1997; Koelsch et al., 2000; Tillmann et al., 2000; Tillmann, 2005; Bigand and Poulin-Charronnat, 2006), providing an example of how unsupervised learning from music might alter representations in the brain. Additionally, behavioral studies of non-industrialized societies who lack much contact with Western culture show pronounced differences from Westerners in many aspects of music perception (McDermott et al., 2016; Jacoby and McDermott, 2017; Jacoby et al., 2019; McPherson et al., 2020), and might plausibly also exhibit differences in the degree or nature of cortical music selectivity. Thus, our data do not show that music selectivity in the brain is independent of experience, but rather that typical exposure to music in Western culture is sufficient for cortical music selectivity to emerge. It remains possible that the brains of people who grow up with less extensive musical exposure than our participants would not display such pronounced music selectivity.

### What does cortical music selectivity represent?

The music-selective component responds strongly to a wide range of music and weakly to virtually all other sounds, demonstrating that it is driven by a set of features that are relatively specific to music. One possibility is that there are simple acoustic features that differentiate music from other types of stimuli. Speech and music are known to differ in their temporal modulation spectra, peaking at 5 Hz and 2 Hz, respectively (Ding et al., 2017), and some theories suggest that these acoustic differences lead to neural specialization for speech vs. music in different cortical regions (Albouy et al., 2020). However, standard auditory models based on spectrotemporal modulation do not capture the perception of speech and music (McDermott and Simoncelli, 2011) or neural responses selective for speech and music (Overath et al., 2015; Kell et al., 2018; Norman-Haignere and McDermott, 2018; Zuk et al., 2020). In particular, the music-selective component responds substantially less to sounds that have been synthesized to have the same spectrotemporal modulation statistics as natural music, suggesting that the music component does not simply represent the audio or modulation frequencies that are prevalent in music (Norman-Haignere and McDermott, 2018).

Our finding that the music-selective component shows high responses to less familiar musical genres places some constraints on what these properties might be, as does the short duration of the stimuli used to characterize music selectivity. For instance, the music-specific features that drive the response are unlikely to be specific to Western music, and must unfold over relatively short timeframes. Features that are common to nearly all music, but not other types of sounds, include stable and sustained pitch organized into discrete note-like elements, and temporal patterning with regular time intervals. Because the music component anatomically overlaps with more general responses to pitch (Norman-Haignere et al., 2015), it is natural to wonder if it represents higher-order aspects of pitch, such as the above-mentioned stability, or discrete jumps from one note to another. However, the high response to drum rhythms in the music component that we observed here indicates that the component is not only sensitive to pitch structure. Instead, this result suggests that melody and rhythm might be jointly analyzed, rather than dissociated, at least at the level of auditory cortex. One possibility is that the underlying neural circuits extract temporally local representations of melody and rhythm motifs that are assembled elsewhere into the representations of contour, key, meter, groove etc. that are the basis of music cognition (Janata et al., 2002; Brett and Grahn, 2007; Lee et al., 2011; Fedorenko et al., 2012; Matthews et al., 2020).

### Limitations

Our paradigm used relatively brief stimuli since music-selective regions are present just outside of primary auditory cortex, where integration periods appear to be short (Overath et al., 2015). And we intentionally used a simple task (intensity discrimination) in order to encourage subjects to attend to all stimuli. But because the responses of auditory cortical neurons have been known to change based on task demands (e.g. Fritz et al. 2003), it is possible that more complex stimuli or tasks would reveal additional aspects of music-selective responses, which might not be present to a similar degree in non-musicians. One relevant point of comparison is the finding that amusic participants with striking pitch perception deficits show pitch-selective auditory cortical responses that are indistinguishable from those of control participants with univariate analyses (Norman-Haignere et al., 2016). Recent evidence suggests it is nonetheless possible to discriminate amusic participants from controls, and to predict participants’ behavioral performance, using fMRI data collected in the context of a pitch task (Albouy et al., 2019). Utilizing a music-related task might produce larger differences between musicians and non-musicians, as might longer music stimuli (compared to the 2-second clips used in this experiment), which could be argued to contain richer melodic, harmonic, and/or rhythmic information.

Finally, our study is limited by the resolution of fMRI. Voxel decomposition is intended to help overcome the spatial limitations of fMRI, and indeed appears to reveal responses that are not evident in raw voxel responses but can be seen with finer-grained measurement substrates such as electrocorticography (Norman-Haignere et al., 2019). But the spatial and temporal resolution of the BOLD signal inevitably constrain what is detectable and place limits on the precision with which we can observe the activity of music-selective neural populations. Music-selective brain responses might well exhibit additional characteristics that would only be evident in fine-grained spatial and temporal response patterns that cannot be resolved with fMRI. Thus, we cannot rule out the possibility that there are additional aspects of music-selective neural responses that might be detectable with other neuroimaging methods (e.g. M/EEG, ECoG) and which are absent or altered in non-musicians.

### Future directions

One of the most interesting open questions raised by our findings is whether cortical music selectivity reflects implicit knowledge gained through typical exposure to music, or whether it is present from birth. These hypotheses could be addressed by testing people with very different musical experiences from non-Western cultures or other populations whose lifetime perceptual experience with music is limited in some way (e.g., people with musical anhedonia, children of deaf adults). It would also be informative to test whether music selectivity is present in infants or young children. Finally, much remains to be learned about the nature of cortical music selectivity, such as what acoustic or musical features might be driving it. The voxel decomposition approach provides one way of answering these questions and exploring the quintessentially human ability for music.

## ACKNOWLEDGEMENTS

This work was supported by NSF grant BCS-1634050 to J.M. and NIH grant DP1HD091947 to N.K. The Athinoula A. Martinos Imaging Center at MIT is supported by the NIH Shared instrumentation grant #S10OD021569. Thank you to the McDermott lab for useful comments on an earlier version of this manuscript.

## SUPPLEMENTAL FIGURES

**Supplemental Figure S1.**
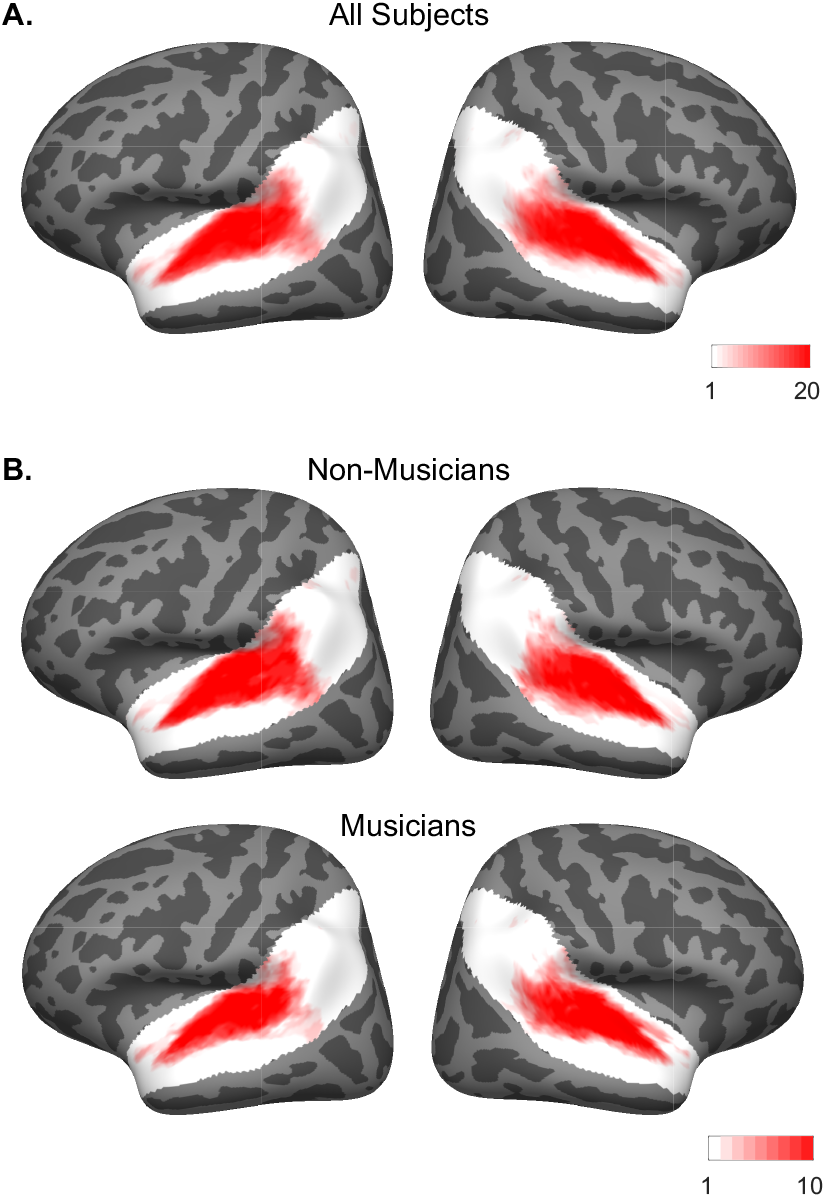
Subject overlap maps showing which voxels were selected in individual subjects to serve as input to the voxel decomposition algorithm. To be selected, a voxel must display a significant (p < 0.001, uncorrected) response to sound (pooling over all sounds compared to silence), and produce a reliable response pattern to the stimuli across scanning sessions (see equations in Methods section). The white area shows the anatomical constraint regions from which voxels were selected. (**A**) Overlap map for all twenty subjects. (**B**) Overlap maps for non-musicians and musicians separately, illustrating that the anatomical location of the selected voxels was largely similar across groups.

**Supplemental Figure S2.**
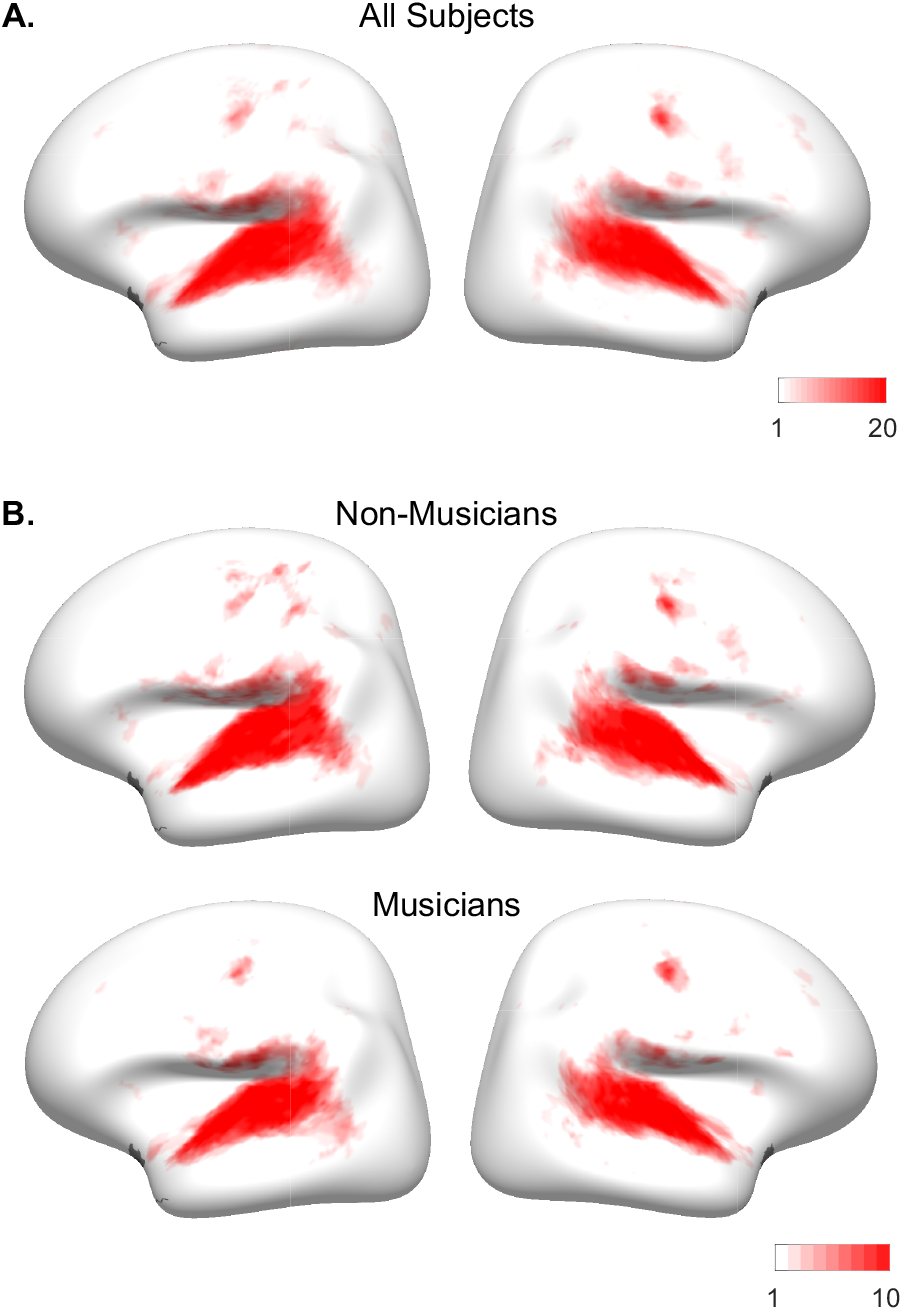
Subject overlap maps showing which voxels pass the selection criteria as described in **Supplemental Figure S1**, but without any anatomical mask applied before selecting voxels.

**Supplemental Figure S3.**
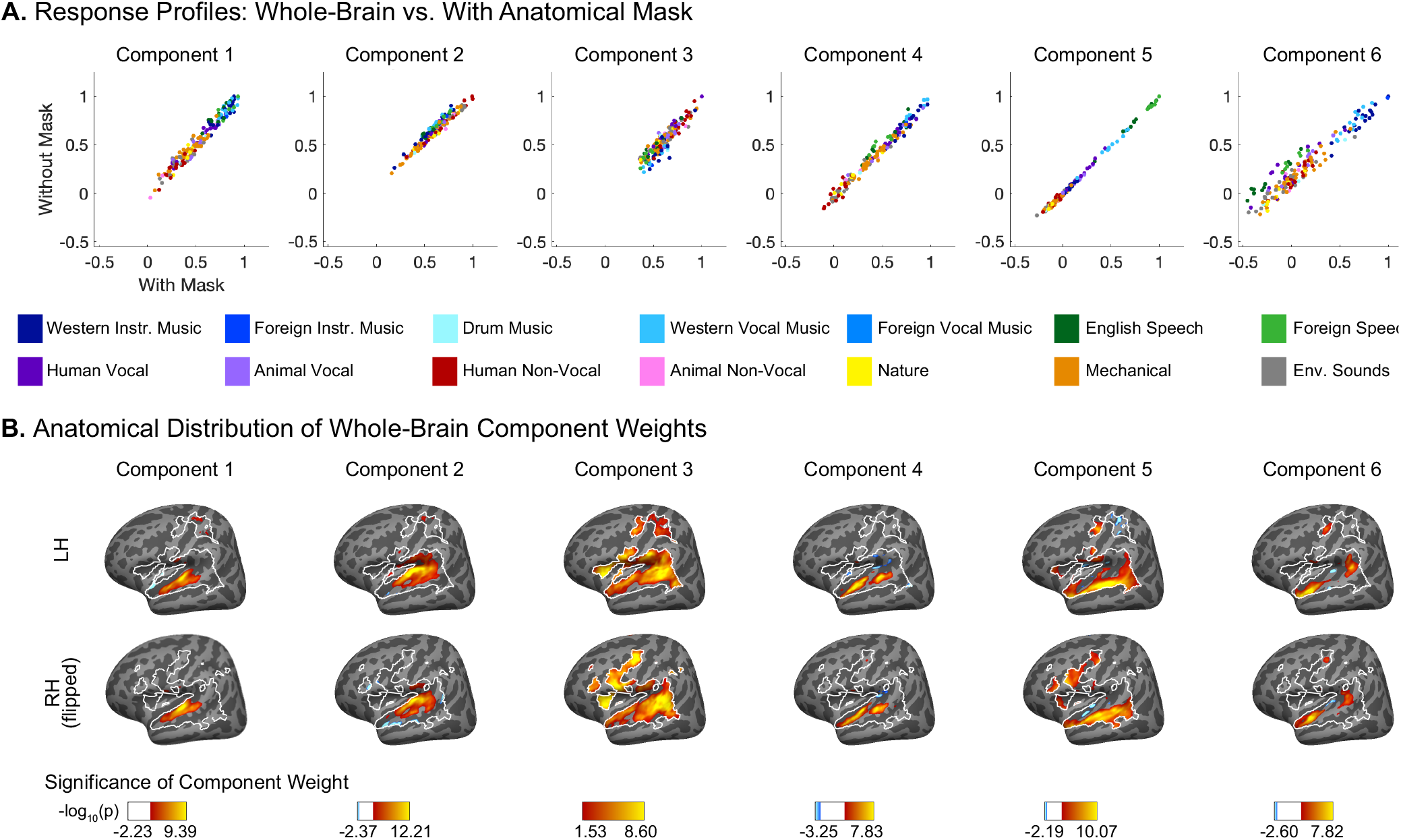
Similarity between components with anatomical mask vs. whole-brain. (**A**) Scatter plots showing the components inferred from all 20 participants, using the voxel decomposition algorithm both with and without the anatomical mask shown in **Supplemental Figure S1**. Individual sounds are colored according to their semantic category. (**B**) Spatial distribution of whole-brain component voxel weights, computed using a random effects analysis of participants’ individual component weights. Weights are compared against 0; p values are logarithmically transformed (-log10[p]). The white outline indicates the voxels that were both sound-responsive (sound vs. silence, p < 0.001 uncorrected) and split-half reliable (r > 0.3) at the group level. The color scale represents voxels that are significant at FDR q = 0.05, with this threshold being computed for each component separately. Voxels that do not survive FDR correction are not colored, and these values appear as white on the color bar. The right hemisphere (bottom row) is flipped to make it easier to visually compare weight distributions across hemispheres.

**Supplemental Figure S4.**
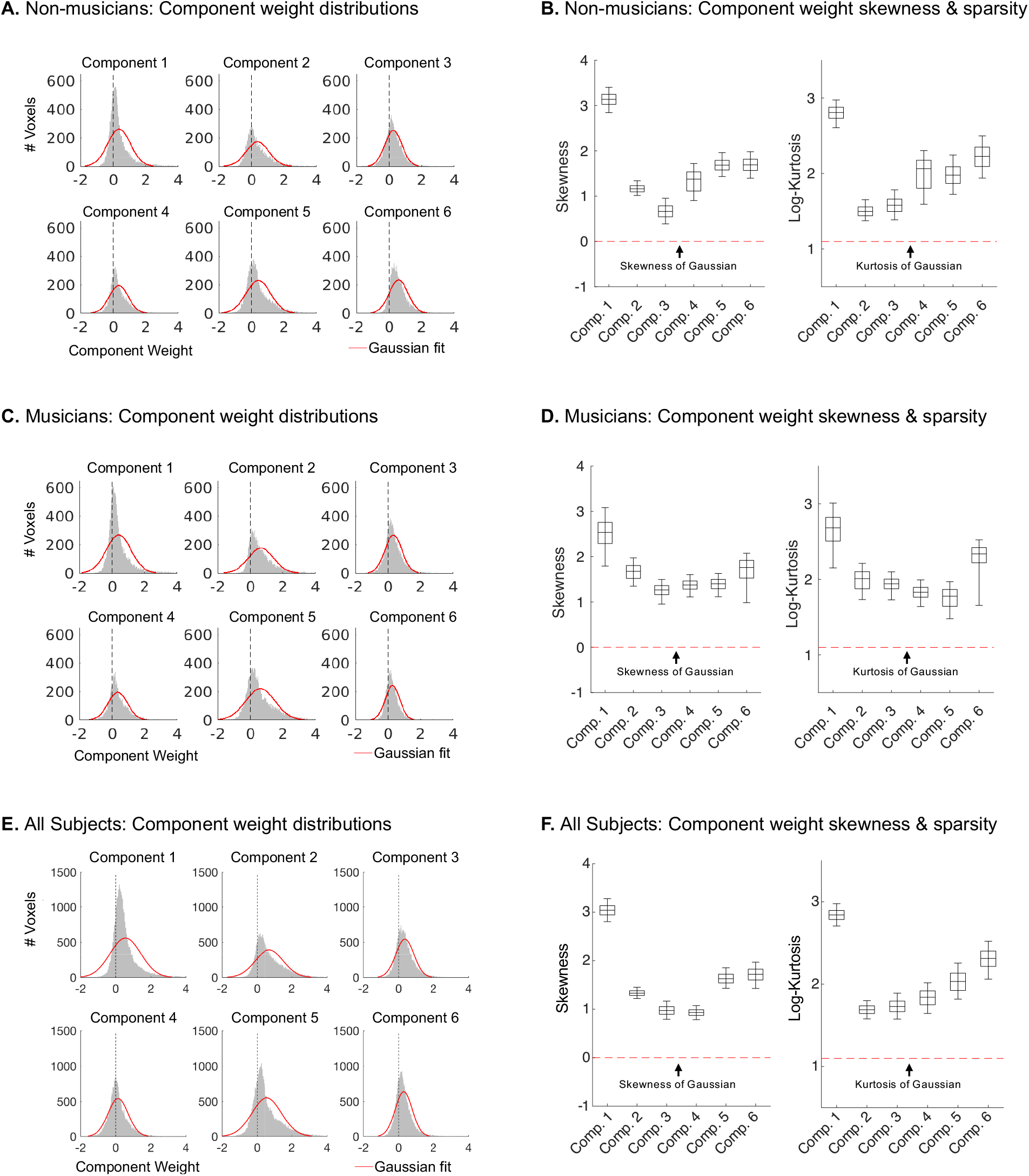
Like ICA, the voxel decomposition method searches amongst the many possible solutions to the factorization problem for components that have a maximally non-Gaussian distribution of weights across voxels. (**A**) Histograms showing the weight distributions for each component inferred from non-musicians (n = 10) can be seen here, along with their Gaussian fits (red). (**B**) Skewness and log-kurtosis (a measure of sparsity) for each component inferred from musicians (n = 10), illustrating that the inferred components are skewed and sparse compared to a Gaussian (red dotted lines). Box-and-whisker plots show central 50% (boxes) and central 95% (whiskers) of the distribution for each statistic (via bootstrapping across subjects). For both the weight distribution histograms and analyses of non-Gaussianity, we used independent data to infer components (runs 1-24) and to measure the statistical properties of the component weights (runs 25-48). (**C, D**) Same as **A** & **B**, but for the components inferred from musicians (n = 10). (**E, F**) Same as **A** & **B**, but for the components inferred from all 20 participants

**Supplemental Figure S5.**
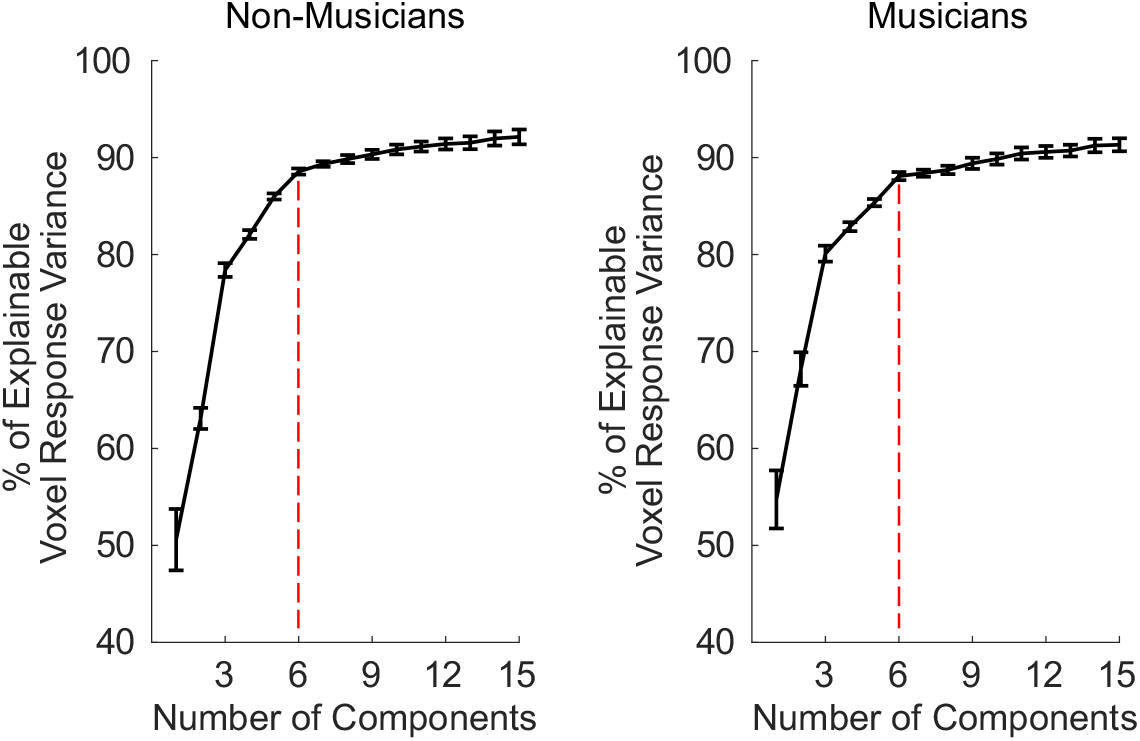
The proportion of voxel response variance explained by different numbers of components, for both non-musicians (left) and musicians (right). The figure plots the median variance explained across voxels (noise corrected by split-half reliability), calculated separately for each subject and then averaged across the ten subjects in each group. Error bars plot one standard error of the mean across subjects. For both groups, 6 components were sufficient to explain over 88% of the noise-corrected variance.

**Supplemental Figure S6.**
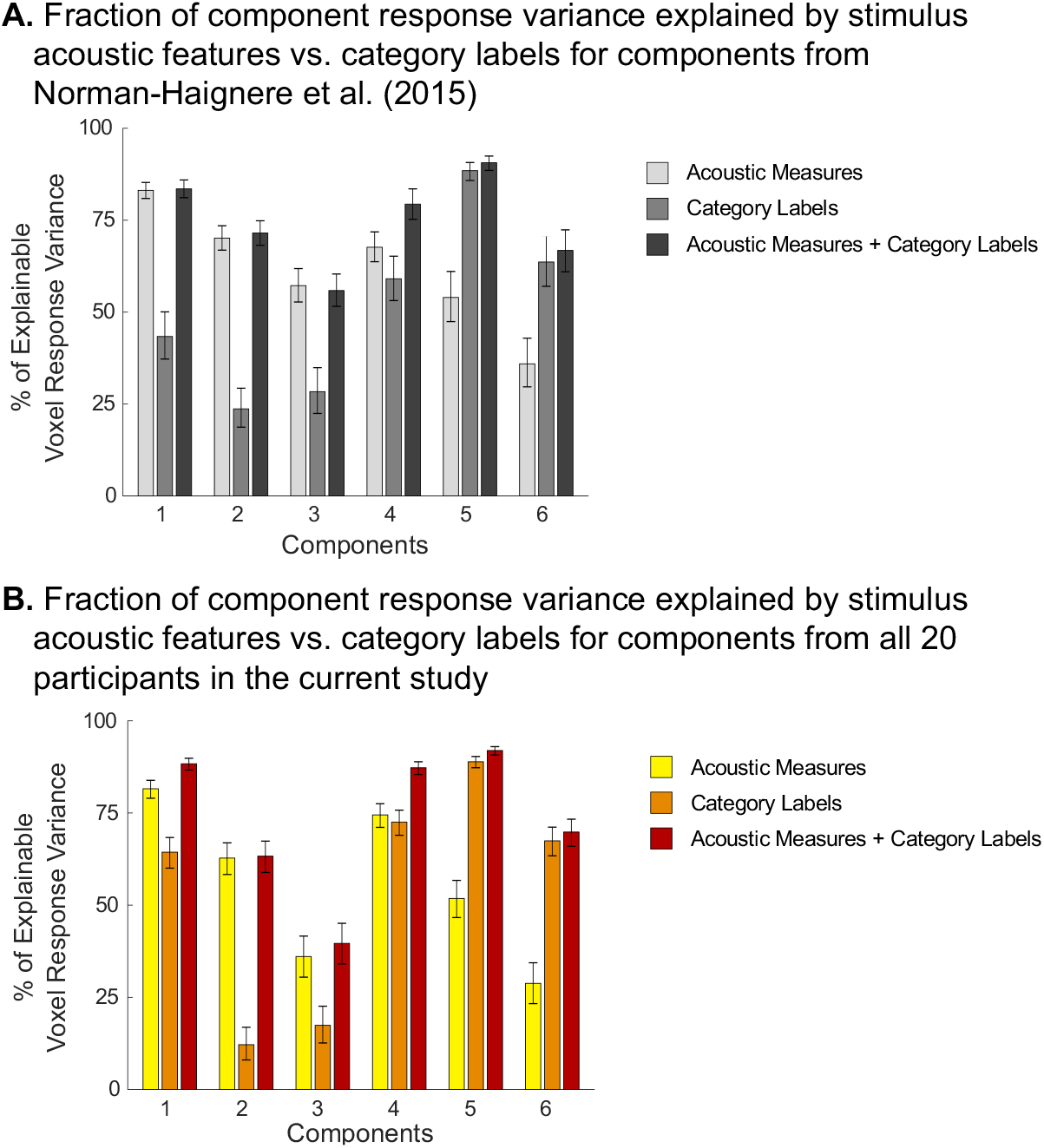
Total amount of component response variation explained by (1) all acoustic measures (frequency content and spectrotemporal modulation energy), (2) all category labels (as assigned by Amazon Mechanical Turk workers), and (3) the combination of acoustic measures and category labels. (**A**) Results for components from previous study (Norman-Haignere et al., 2015). For components 1–4, category labels explained little additional variance beyond that explained by acoustic features. For Components 5 (speech-selective) and 6 (music-selective), category labels explained most of the response variance, and acoustic features accounted for little additional variance. (**B**) Same as **A** but for the components inferred from all 20 participants in the current study.

**Supplemental Figure S7.**
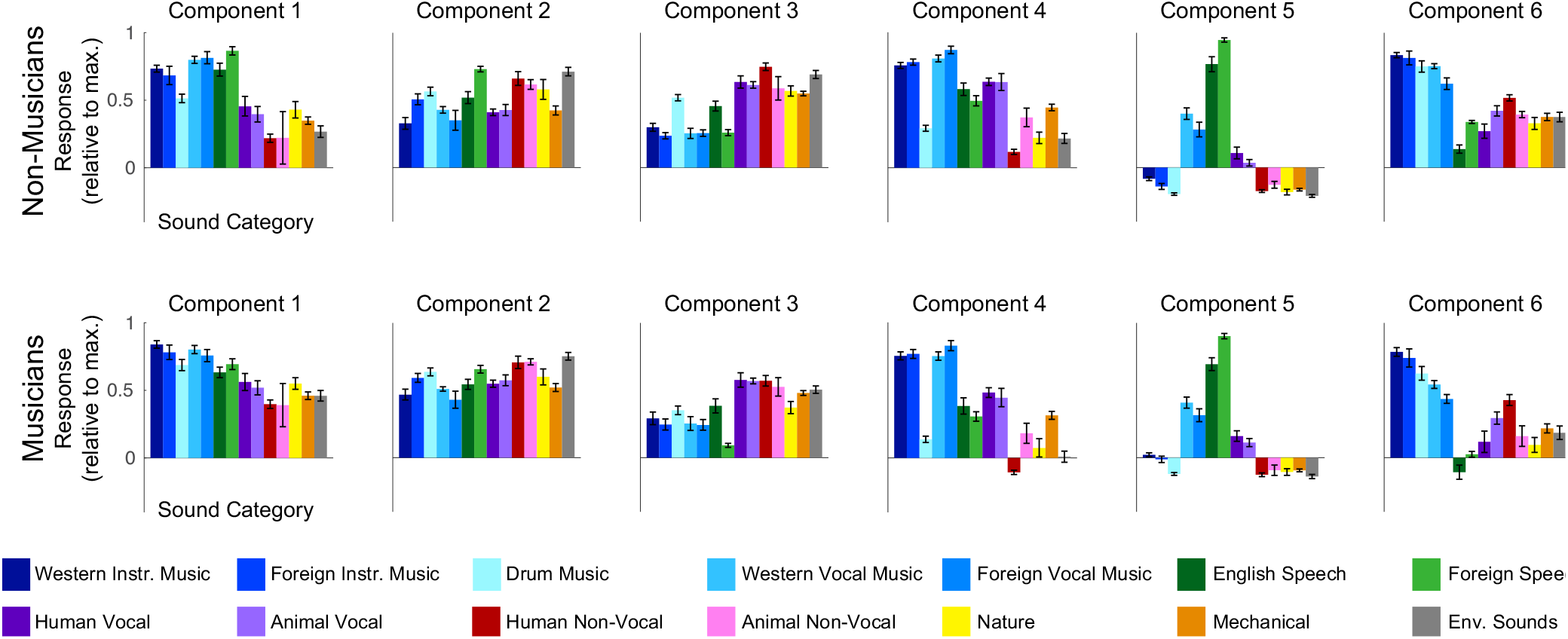
Response profiles discovered using the probabilistic non-negative matrix factorization (NMF) method (see details in **Supplemental Information**). These components were very highly correlated with those inferred using the ICA-based voxel decomposition method presented in the main text, with the main difference between the two methods being the mean response profile magnitude (i.e. the “offset” from baseline). Because this mean response varies depending on the details of the analysis used to infer the components, while the components themselves remain highly similar, we chose to quantify selectivity using a measure (Cohen’s d) that does not take the baseline into account but rather quantifies the separation between stimulus categories within the response profile.

**Supplemental Figure S8.**
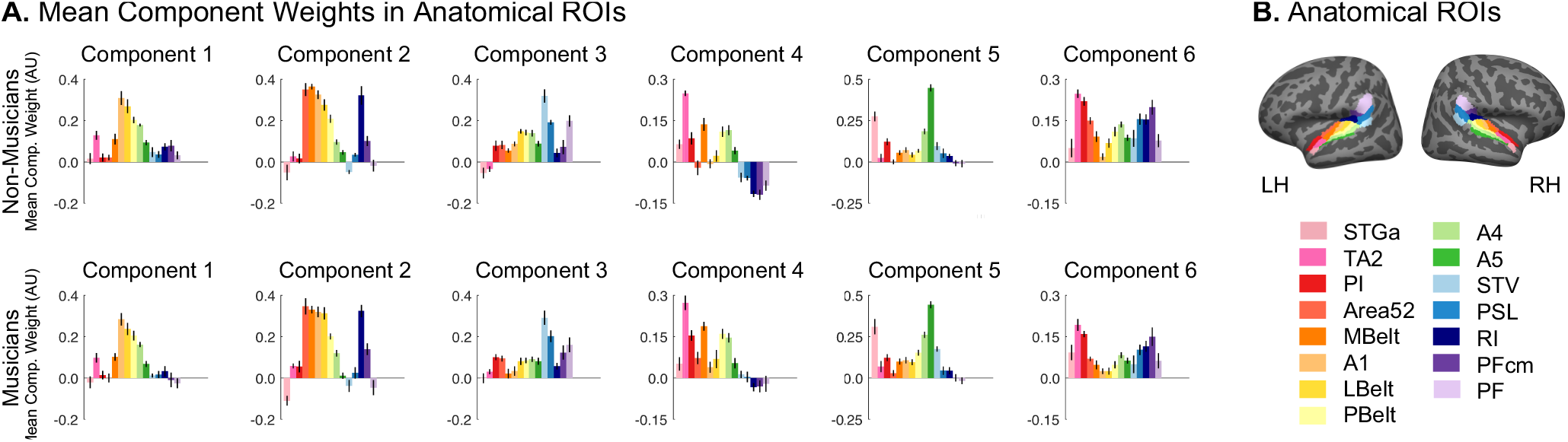
Mean component voxel weights within standardized anatomical parcels. (**A**) Mean component weight in a set of fifteen anatomical parcels from Glasser et al. (2016), plotted separately for non-musicians (top) and musicians (bottom). Error bars plot one standard error of the mean across participants. (**B**) Selected anatomical parcels from Glasser et al. (2016), chosen to fully encompass the superior temporal plane and superior temporal gyrus (STG).

**Supplemental Figure S9.**
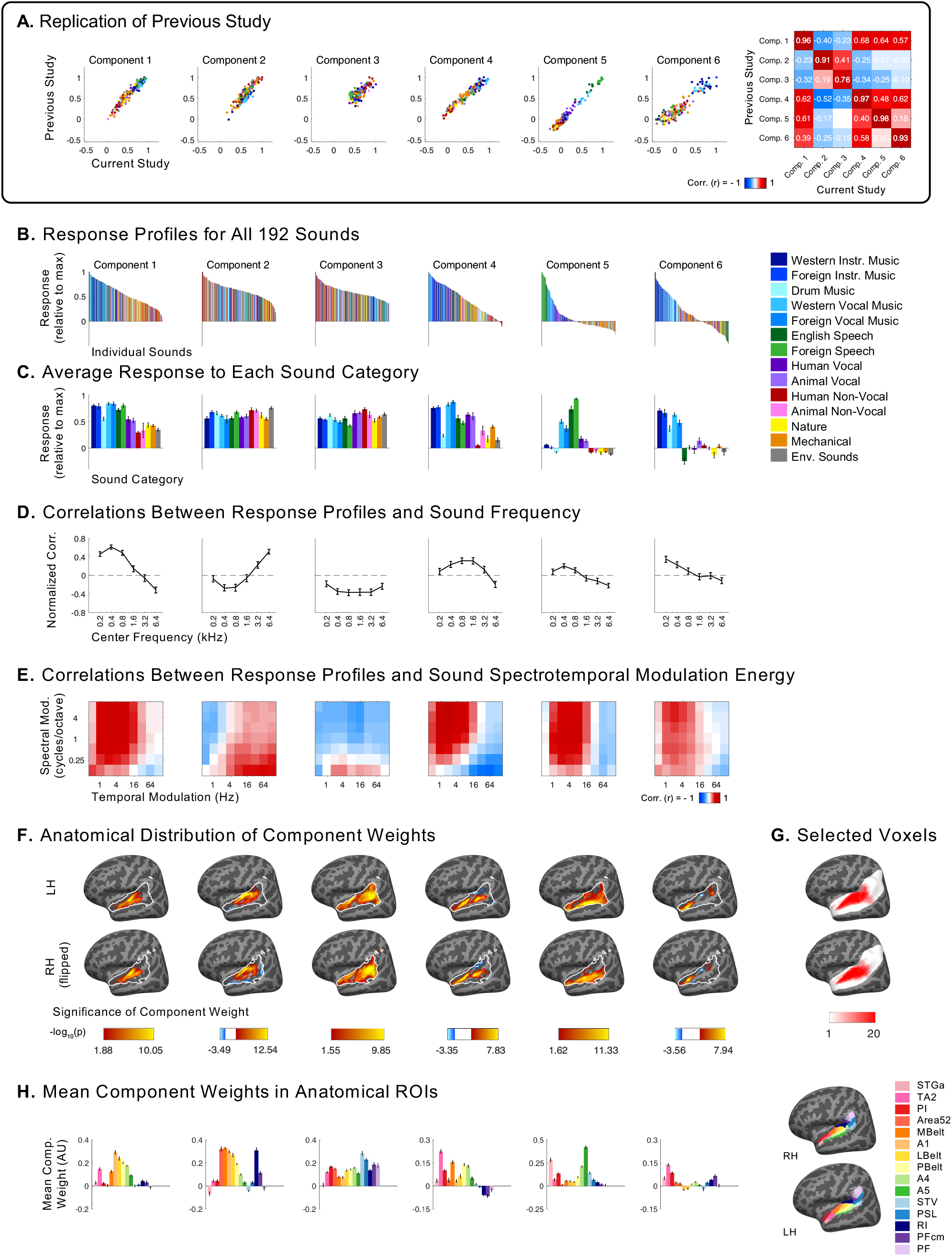
Independent components inferred from voxel decomposition of auditory cortex of all 20 participants (as compared to the components in **Figures 2–5**, which were inferred from musicians and non-musicians separately). Additional plots are included here to show the extent of the replication of the results of Norman-Haignere et al. (2015). (**A**) Scatterplots showing the correspondence between the components from our previous study (y-axis) and those from the current study (x-axis). Only the 165 sounds that were common between the two studies are plotted. Sounds are colored according to their semantic category, as determined by raters on Amazon Mechanical Turk. (**B**) Response profiles of components inferred from all participants (n = 20), showing the full distribution of all 192 sounds. Sounds are colored according to their category. Note that “Western Vocal Music” stimuli were sung in English. (**C**) The same response profiles as above, but showing the average response to each sound category. Error bars plot one standard error of the mean across sounds from a category, computed using bootstrapping (10,000 samples). (**D**) Correlation of component response profiles with stimulus energy in different frequency bands. (**E**) Correlation of component response profiles with spectrotemporal modulation energy in the cochleograms for each sound. (**F**) Spatial distribution of component voxel weights, computed using a random effects analysis of participants’ individual component weights. Weights are compared against 0; p values are logarithmically transformed (-log10[p]). The white outline indicates the 2,249 voxels that were both sound-responsive (sound vs. silence, p < 0.001 uncorrected) and split-half reliable (r > 0.3) at the group level. The color scale represents voxels that are significant at FDR q = 0.05, with this threshold being computed for each component separately. Voxels that do not survive FDR correction are not colored, and these values appear as white on the color bar. The right hemisphere (bottom row) is flipped to make it easier to visually compare weight distributions across hemispheres. (**G**) Subject overlap maps showing which voxels were selected in individual subjects to serve as input to the voxel decomposition algorithm (same as **Supplemental Figure S1A**). To be selected, a voxel must display a significant (p < 0.001, uncorrected) response to sound (pooling over all sounds compared to silence), and produce a reliable response pattern to the stimuli across scanning sessions (see equations in Methods section). The white area shows the anatomical constraint regions from which voxels were selected. (**H**) Mean component voxel weights within standardized anatomical parcels from Glasser et al. (2016), chosen to fully encompass the superior temporal plane and superior temporal gyrus (STG). Error bars plot one standard error of the mean across participants.

## SUPPLEMENTAL INFORMATION

### Behavioral data acquisition & analysis

To validate participants’ self-reported musicianship, we measured participants’ abilities on a variety of psychoacoustical tasks for which prior evidence suggested that musicians would outperform non-musicians. For all psychoacoustic tasks, stimuli were presented using Psychtoolbox for Matlab (Brainard, 1997). Sounds were presented to participants at 70dB SPL over circumaural Sennheiser HD280 headphones in a soundproof booth (Industrial Acoustics) (SPL level was computed without any weighting across time or frequency). After each trial, participants were given feedback about whether or not they had answered correctly. Group differences for each task were measured using non-parametric Wilcoxon rank sum tests.

#### Pure tone frequency discrimination

Because musicians have superior frequency discrimination abilities when compared to non-musicians (Spiegel and Watson, 1984; Kishon-Rabin et al., 2001; Micheyl et al., 2006), we first measured participants’ pure tone frequency discrimination thresholds using an adaptive two-alternative forced choice (2AFC) task. In each trial, participants heard two pairs of tones. One of the tone pairs consisted of two identical 1 kHz tones, while the other tone pair contained a 1 kHz tone and a second tone of a different frequency. Participants determined which tone interval contained the frequency change. The magnitude of the frequency difference was varied adaptively using a 1-up 3-down procedure (Levitt, 1971), which targets participants’ 79.4% threshold. The frequency difference was changed initially by a factor of two, which was reduced to a factor of √2 after the fourth reversal. Once 10 reversals had been measured, participants’ thresholds were estimated as the average of these 10 values. Multiple threshold estimations were obtained per participant (3 threshold estimations for the first seven participants, and 5 for the remaining 13 participants), and then averaged.

#### Synchronized tapping to an isochronous beat

Sensorimotor abilities are crucial to musicianship, and finger tapping tasks show some of the most reliable effects of musicianship (Repp, 2005, 2010; Bailey and Penhune, 2010). Participants were asked to tap along with an isochronous click track. They heard ten 30-second click blocks, separated by 5 seconds of silence. The blocks varied widely in tempo, with interstimulus intervals ranging from 200ms to 1 second (tempos of 60 to 300 bpm). Each tempo was presented twice, and the order of tempi was permuted across participants. We recorded the timing of participants’ responses using a tapping sensor we constructed and have used in previous studies (Jacoby and McDermott, 2017; Polak et al., 2018). We then calculated the difference between participants’ response onsets and the actual stimulus onsets. The standard deviation of these asynchronies between corresponding stimulus and response onsets was used as a measure of sensorimotor synchronization ability (Polak et al., 2018).

#### Melody discrimination

Musicians have also been reported to outperform non-musicians on measures of melodic contour and interval discrimination (Fujioka et al., 2004; McDermott et al., 2010; McPherson and McDermott, 2018). In each trial, participants heard two five-note melodies, and were asked to judge whether the two melodies were the same or different. Melodies were composed of notes that were randomly drawn from a log uniform distribution of semitone steps from 150Hz to 270Hz. The second melody was transposed up by half an octave and was either identical to the first melody or contained a single note had that had been altered either up or down by 1 or 2 semitones. Half of the trials contained a second melody that was the same as the first melody, while 25% contained a pitch change that preserved the melodic contour and the remaining 25% contained a pitch change that violated the melodic contour. There were 20 trials per condition (same/different melody x same/different contour x 1/2 semitone change), for a total of 160 trials. This task was modified from (McPherson and McDermott, 2018).

#### “Sour note” detection

To measure participants’ knowledge of Western music, we also measured participants’ ability to determine whether a melody conforms to the rules of Western music theory. The melodies used in this experiment were randomly generated from a probabilistic generative model of Western tonal melodies that creates a melody on a note-by-note basis according to the principles that (1) melodies tend to be limited to a narrow pitch range, (2) note-to-note intervals tend to be small, and (3) the notes within the melody conform to a single key (Temperley, 2008). In each trial of this task, participants heard a 16-note melody and were asked to determine whether the melody contained an out-of-key (“sour”) note. In half of the trials, one of the notes in the melody was modified so that it was rendered out of key. The modified notes were always scale degrees 1, 3, or 5 and they were modified by either 1 or 2 semitones accordingly so that they were out of key (i.e. scale degrees 1 and 5 were modified by 1 semitone, and scale degree 3 was modified by 2 semitones). Participants judged whether the melody contained a sour note (explained as a “mistake in the melody”). There were 20 trials per condition (modified or not x 3 scale degrees), for a total of 120 trials. This task was modified from (McPherson and McDermott, 2018).

### Psychoacoustic results

As predicted, musicians outperformed non-musicians on all behavioral psychoacoustic tasks, replicating prior findings (**Supplemental Figure S10**). Consistent with previous reports (Spiegel and Watson, 1984; Kishon-Rabin et al., 2001; Micheyl et al., 2006), musicians performed slightly better on the frequency discrimination task (median discrimination threshold = 0.51%, SD = 0.12%) than non-musicians (median discrimination threshold = 0.57%, SD = 0.23%); this difference was marginally significant (z = −1.32, p = 0.09, effect size r = −0.30, one-tailed Wilcoxon rank sum test, **Supplemental Figure S10A**). Musicians were also better able to synchronize their finger tapping with an isochronous beat, showing significantly less variability than non-musicians in their response (SD = 22.4ms) than non-musicians (SD = 37.7ms, z = −2.68, p < 0.01, effect size r = −0.60, one-tailed Wilcoxon rank sum test, **Supplemental Figure S10B**). When presented with musical melodies, musicians were better able to discriminate between two similar melodies (musician median ROC area = 0.82, SD = 0.08, non-musician mean ROC area = 0.66, SD = 0.09, z = 3.21, p < 0.001, effect size r = 0.72, one-tailed Wilcoxon rank sum test, **Supplemental Figure S10C**), and to detect scale violations within melodies (musician median ROC area = 0.89, SD = 0.06, non-musicians median ROC area = 0.70, SD = 0.10, z = 3.44, p < 0.001, effect size r = 0.77, one-tailed Wilcoxon rank sum test, **Supplemental Figure S10D**). These behavioral effects validate our participants’ self-reported status as trained musicians or non-musicians.

**Supplemental Figure S10.**
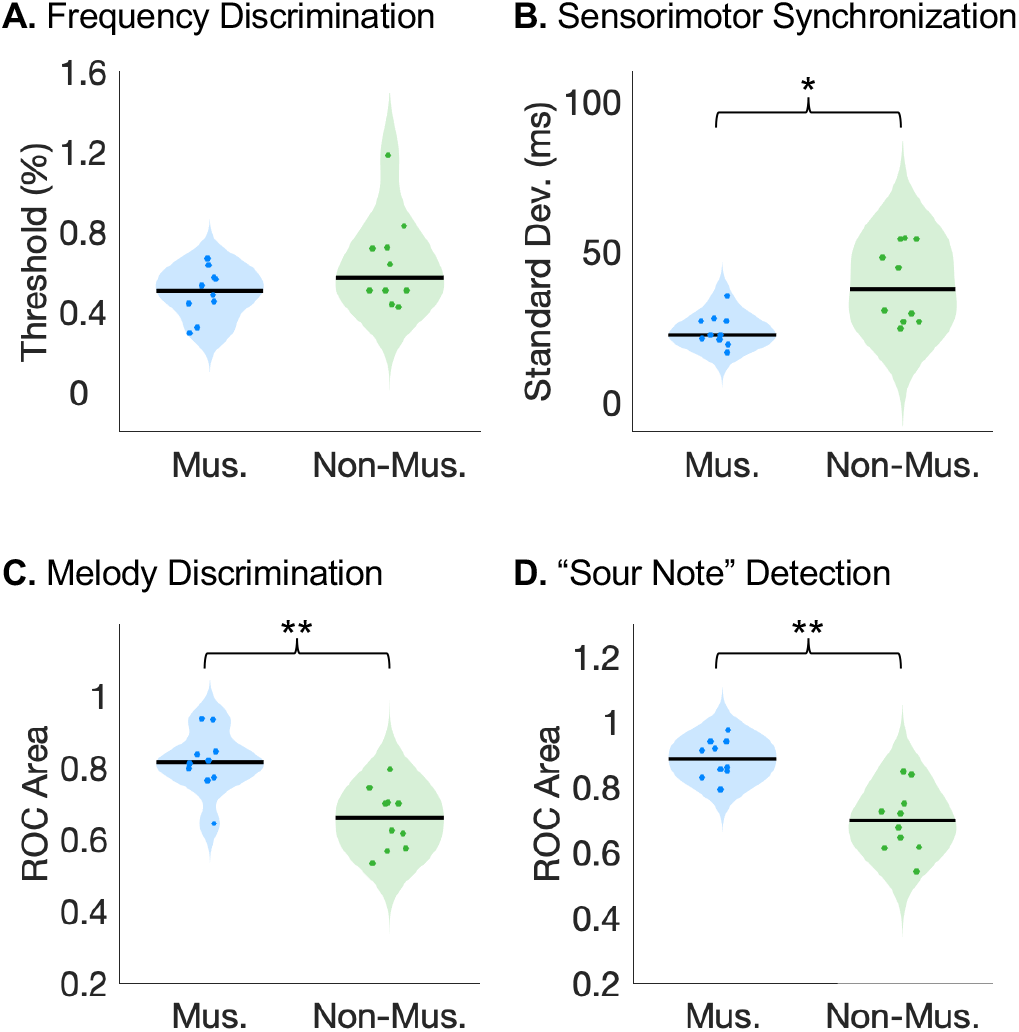
Musicians outperform non-musicians on psychoacoustic tasks. (**A**) Participants’ pure tone frequency discrimination thresholds were measured using a 1-up 3-down adaptive two-alternative forced choice (2AFC) task, in which participants indicated which of two pairs of tones were different in frequency. Note that lower thresholds correspond to better performance. (**B**) Sensorimotor synchronization abilities were measured by instructing participants to tap along with an isochronous beat at various tempos, and comparing the standard deviation of the difference between participants’ response onsets and the actual stimulus onsets. (**C**) Melody discrimination was measured using a 2AFC task, in which participants heard two five-note melodies (with the second one transposed up by a tritone) and were asked to judge whether the two melodies were the same or different. (**D**) We measured participants’ ability to determine whether a melody conforms to the rules of Western music theory by creating 16-note melodies using a probabilistic generative model of Western tonal melodies (Temperley, 2008), and instructing participants to determine whether or not the melody contained an out-of-key (“sour”) note. Colored dots represent individual participants, and the median for each participant group is indicated by the horizontal black line. Mus. = musicians, Non-Mus. = non-musicians. * = significant at p < 0.01 one-tailed, ** = significant at p < 0.001 one-tailed.

### Replication of Norman-Haignere et al. (2015) using data from all 20 participants

In addition to conducting the voxel decomposition analysis separately for musicians and non-musicians, we were able replicate the full results from Norman-Haignere et al. (2015) using data from all 20 participants from both groups. The components from this combined analysis were used when directly comparing component weights between groups (described in Results sections titled “Music component weight magnitude differs only slightly between musicians and non-musicians” and “Similar anatomical distribution of music component weights in musicians and non-musicians”). Here, we describe that analysis in more detail, and explain more about the four components that are selective for acoustic stimulus features.

In our previous study (Norman-Haignere et al., 2015), as in the analyses described above in the current study, prior to applying the voxel decomposition algorithm, each participant’s responses were de-meaned across voxels (see Methods), such that each participant had the same mean response (across voxels) for a given sound. This normalization was included to prevent the voxel decomposition algorithm from discovering additional components that were driven by a single participant (e.g. due to non-replicable sources of noise, such as motion during a scan). However, this analysis step would also remove any group difference in the average response to certain sounds (e.g. music stimuli). To prevent this effect from removing differences between musicians and non-musicians that might be of interest, we ran the voxel decomposition algorithm without demeaning by individual participants. When we varied the number of components as we normally do in order to determine the number of components to use for ICA, we found that the best results were obtained with 8 components, which included the expected set of 6 plus two “extra” components that emerged as a result of the omitted normalization step. We know this to be the case, because if we include the normalization step, we find the expected set of 6 components.

Of the 8 components derived from the non-demeaned data, six of them were each very similar to one of the 6 components from Norman-Haignere et al. (2015) and accounted for 87.54% of voxel response variance. Because the components inferred using ICA have no order, we first used the Hungarian algorithm (Kuhn, 1955) to optimally reorder the components, maximizing their correlation with the components from our previous study. As expected, the reordered components were highly correlated, with r-values ranging from 0.76 to 0.98 (**Supplemental Figure S9A**; see **Supplemental Figure S9B** for the response profiles of the components, and **Supplemental Figure S9C** for the profiles averaged within sound categories). To confirm that these strong correlations are not simply an artifact of the Hungarian algorithm matching procedure, we ran a permutation test in which we re-ordered the sounds within each component 1,000 times, each time using the Hungarian algorithm to match these permuted components with those from our previous study. The resulting correlations between the original components their corresponding permuted components were very low (mean r-values ranging from 0.086 to 0.098), with the maximum correlation over all 1,000 permutations not exceeding r = 0.3 for any component.

The additional two components were much less correlated with any of the six original components, with the strongest correlation being r = 0.28. As expected, the weights of these additional two components were concentrated in a small number of participants (one almost entirely loading onto a single non-musician participant, and the other onto a small group comprised of both musicians and non-musicians). For this reason, we omitted these two components from further analyses and focused on the set of six components that closely match those discussed previously (**Supplemental Figure S9B-H**). The non-Gaussianity of these 6 components can be seen in **Supplemental Figure S4A** (skewness ranging from 1.06 to 2.96, log-kurtosis ranging from 1.70 to 2.79).

As in Norman-Haignere et al. (2015), four of the components were selective for different acoustic properties of sound (**Supplemental Figure S9D** & **E**), while two components were selective for speech (component 5) and music respectively (component 6) (**Supplemental Figure S9B** & **C)**. The components replicated all of the functional and anatomical properties from our prior study, which we briefly describe here.

Components 1 and 2 exhibited high correlations between their response profiles and measures of stimulus energy in either low (component 1) or high frequency bands (component 2) (**Supplemental Figure S9D**). The group anatomical weights for components 1 and 2 concentrated in the low and high-frequency regions of primary auditory cortex (**Supplemental Figure S9F** & **H**) (Rauschecker et al., 1995; Humphries et al., 2010; Da Costa et al., 2011; Baumann et al., 2013). We did not measure tonotopy in the individual participants from this study, but our previous study did so, and found a close correspondence between individual participant tonotopic maps and the weights for these two components. Components also showed tuning to spectrotemporal modulations (**Supplemental Figure S9E**), with a tradeoff between selectivity for fine spectral and slow temporal modulation (components 1 and 4) verses coarse spectral and fast temporal modulation (components 2 and 3) (Singh and Theunissen, 2003; Rodríguez et al., 2010). Component 4, which exhibited selectivity for fine spectral modulation, was concentrated anterior to Heschl’s gyrus (component 4, **Supplemental Figure S9F** & **H**), similar to prior work that has identified tone-selective regions in anterolateral auditory cortex in humans (Patterson et al., 2002; Penagos et al., 2004; Norman-Haignere et al., 2013). Conversely, selectivity for coarse spectral modulation and fast temporal modulation was concentrated in posterior regions of auditory cortex (component 3, **Supplemental Figure S9F** & **H**) (Santoro et al., 2014), consistent with previous studies reporting selectivity for sound onsets in caudal areas of human auditory cortex (Hamilton et al., 2018).

The two remaining components responded selectively to speech and music, respectively (component 5 and 6, **Supplemental Figure S9C**), and were not well accounted for using acoustic properties alone (**Supplemental Figure S9B**). The weights for the speech-selective component (component 5) were concentrated in the middle portion of the superior temporal gyrus (midSTG, **Supplemental Figure S9F** & **H**), as expected (Scott et al., 2000; Hickok and Poeppel, 2007; Overath et al., 2015). In contrast, the weights for the music-selective component (component 6) were most prominent anterior to PAC in the planum polare, with a secondary cluster posterior to PAC in the planum temporale (**Supplemental Figure S9F** & **H**) (Ohnishi et al., 2001; Margulis et al., 2009; Dick et al., 2011; Fedorenko et al., 2012; Angulo-Perkins et al., 2014; Armony et al., 2015; Norman-Haignere et al., 2015).

These results closely replicate the functional organization of human auditory cortex reported by Norman-Haignere et al. (2015), including the existence and anatomical location of inferred music-selective neural populations.

### Direct group comparisons of music selectivity

#### Music component selectivity

One natural extension of the analyses presented in this paper is to directly compare music selectivity in our group of non-musicians to that observed in our group of musicians. To compare the selectivity of the music components inferred from each group, we computed Cohen’s *d* between the distribution of component responses to music stimuli (“Western instrumental,” “Non-Western instrumental,” “Western vocal,” “Non-Western vocal,” and “drums”) and the distribution of component responses to non-music stimuli (all other sound categories).

The significance of the observed group difference was evaluated using a nonparametric test in which we permuted participant groupings (i.e. randomly assigning 10 participants to group A, and the remaining 10 to group B), and inferred a set of components for each permuted group. We then calculated Cohen’s d for each group’s music component and computed the absolute value of the difference between these two values. We did this 1000 times to build up a null distribution of Cohen’s d differences, and then compared this to the observed difference in Cohen’s d for non-musicians vs. musicians and found the observed difference to be non-significant (p = 0.21, one-tailed).

However, it was not possible to conduct a power analysis using the data from Norman-Haignere et al. (2015) to determine how large of a difference we are powered to detect with our sample size, because the component analysis is sometimes unstable when the number of unique participants drops below 10, which made it impossible to bootstrap and infer a large number of sets of components. For that reason, we cannot rule out the possibility that a small difference does exist, but that this test is underpowered.

#### Music component weight magnitude & power analysis

We thought it would be interesting to directly compare the magnitude of the weights of the music-selective component between expert musicians and non-musicians. So that individual participants’ component weight magnitudes could be compared in a meaningful way, we planned to run the voxel decomposition analysis on the data from all 20 participants (as reported above), and directly compare the weights of musicians vs. non-musicians for the resulting music component. Although there are many ways to summarize a participant’s component magnitude, we used the median weight over their voxels with the top 10% of component weights (using independent data to select the voxels vs. quantify selectivity), though results were robust to the fraction of voxels selected (i.e. measuring participants’ median weight over the top 5%, 7.5%, 10%, 15%, and 20% of voxels led to similar results). This decision to select a subset of voxels was made because music selectivity is typically sparse and limited to a small fraction of voxels, and we thought it reasonable to expect the largest group difference in the regions of auditory cortex with the highest music component weights.

To get a sense for how large a group difference we would be able to reliably detect given our sample size, we conducted a power analysis using the data from Norman-Haignere et al. (2015). We compared the music component weights for the participants in that study (n = 10) with a second population of 10 participants created by sampling participants with replacement and then shifting their component weights by various amounts (ranging from 0% to 100% in increments of 5%), representing various models for how the music component weights might change in musicians. The difference between the groups’ median weights was computed, and the significance of this group difference was assessed by permuting participant groupings 1,000 times. For each shift amount, we repeated this entire procedure 1,000 times, each time sampling a new set of 10 participants for each group. The probability of detecting a significant group difference for each shift amount was recorded, and the results showed that we were able to detect a significant group difference 80% of the time only when the two groups’ median weights differed by 47%.

This power analysis suggests that with our sample size, we are only able to detect a relatively large difference in music component weight magnitude between the groups. With this in mind, we performed this analysis and found the strength of the music component was slightly higher in musicians compared with non-musicians, but this difference did not reach significance (p = 0.11, two-tailed nonparametric test permuting subject groupings 10,000 times). A Bayesian independent-sample t-test was inconclusive (t(18) = 1.68, p = 0.11, BF_10_ = 1.03; prior on the effect size following a Cauchy distribution with 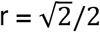), which suggests that the data are equally likely under the null hypothesis that groups do not differ in their music component weight magnitude vs. the alternative hypothesis that they do. Together, these results do not rule out differences between the strength of music selectivity in individuals at the two extremes of musical training, but they suggest that any such differences are not large.

As explained above, we thought that restricting this analysis to the most music-selective voxels (i.e. the voxels with the highest music component weights) would be most likely to detect any group difference in weight magnitude. However, we also tried a simpler analysis in which we ran an ROI x hemisphere repeated-measures ANOVA on participants’ weights for the component inferred from all 20 participants, and including “group” as a between-subjects factor. When we do this, we still find a significant main effect of ROI (F(3,54) = 37.89, p = 2.56e-13), no significant main effect hemisphere (F(1,18) = 0.84, p = 0.37), as was found in the corresponding analyses within each group that are reported in the main text. However, we also found no significant main effect of group (F(1,18) = 2.81, p = 0.11), or any significant 2-way or 3-way interactions with group (all p’s > 0.05). Moreover, a Bayesian version of this analysis provides no evidence either for or against an effect of group (BF_inc_ = 1.11). This result is consistent with the power analysis reported above, indicating that while we find no evidence of a difference between musicians’ and non-musicians’ music component weights, we do not have enough statistical power to rule out the possibility that a small difference does exist.

### Parametric matrix factorization method

In addition to the non-parametric matrix factorization method reported throughout this paper, we repeated our analyses using a probabilistic parametric algorithm also developed and reported in Norman-Haignere et al. (2015), which did not constrain voxel weights to be uncorrelated. This parametric model assumed a skewed and sparse non-Gaussian prior (the Gamma distribution) on the distribution of voxel weights, which constrained them to be positive (unlike the non-parametric method). Because the components discovered using the non-parametric method showed different degrees of skewness and sparsity, the exact shape of the Gamma distribution prior was allowed to vary between components in this parametric analysis. Components were discovered by searching for response profiles and shape parameters that maximized the likelihood of the data, integrating across all possible voxel weights.

Due to the stochastic nature of the model optimization procedure (see Norman-Haignere et al., 2015 for details), the optimization procedure was repeated 25 times each for musicians and non-musicians. For each group, we chose the set of component response profiles with the highest estimated log-likelihood (though all 25 iterations produced very similar results), and used the Hungarian algorithm to match them with the components inferred using the non-parametric method. The sets of components inferred with these two different methods were very highly correlated, with r-values ranging from 0.83 to 0.998 for non-musicians, and from 0.90 to 0.99 for musicians. However, the components did differ somewhat in the mean magnitude of the response profiles (see **Supplemental Figure S7**), plausibly due to the positivity constraint on the component voxel weights.

